# Rational Design of TDP-43 Derived α-Helical Peptide Inhibitors: an *In-Silico* Strategy to Prevent TDP-43 Aggregation in Neurodegenerative Disorders

**DOI:** 10.1101/2023.10.26.564235

**Authors:** Muthu Raj Salaikumaran, Pallavi P. Gopal

## Abstract

TDP-43, an essential RNA/DNA-binding protein, is central to the pathology of neurodegenerative diseases such as Amyotrophic Lateral Sclerosis and Frontotemporal Dementia. Pathological mislocalization and aggregation of TDP-43 disrupts RNA splicing, mRNA stability, and mRNA transport, thereby impairing neuronal function and survival. The formation of amyloid-like TDP-43 filaments is largely facilitated by the destabilization of an α-helical segment within the disordered C-terminal region. In this study, we hypothesized that preventing the destabilization of the α-helical domain could potentially halt the growth of these pathological filaments. To explore this, we utilized a range of *in-silico* techniques to design and evaluate peptide-based therapeutics. Various pathological TDP-43 amyloid-like filament crystal structures were selected for their potential to inhibit the binding of additional TDP-43 monomers to the growing filaments. Our computational approaches included biophysical and secondary structure property prediction, molecular docking, 3D structure prediction, and molecular dynamics simulations. Through these techniques, we were able to assess the structure, stability, and binding affinity of these peptides in relation to pathological TDP-43 filaments. The results of our *in-silico* analyses identified a selection of promising peptides, which displayed a stable α-helical structure, exhibited an increased number of intramolecular hydrogen bonds within the helical domain, and demonstrated high binding affinities for pathological TDP-43 amyloid-like filaments. Additionally, molecular dynamics simulations provided further support for the stability of these peptides, as they exhibited lower root mean square deviations in their helical propensity over 100ns. These findings establish α-helical propensity peptides as potential lead molecules for the development of novel therapeutics against TDP-43 aggregation. This structure-based computational approach for rational design of peptide inhibitors opens a new direction in the search for effective interventions for ALS, FTD, and other related neurodegenerative diseases. The peptides identified as the most promising candidates in this study are currently subject to further testing and validation through both in vitro and in vivo experiments.

**Graphical Abstract:** 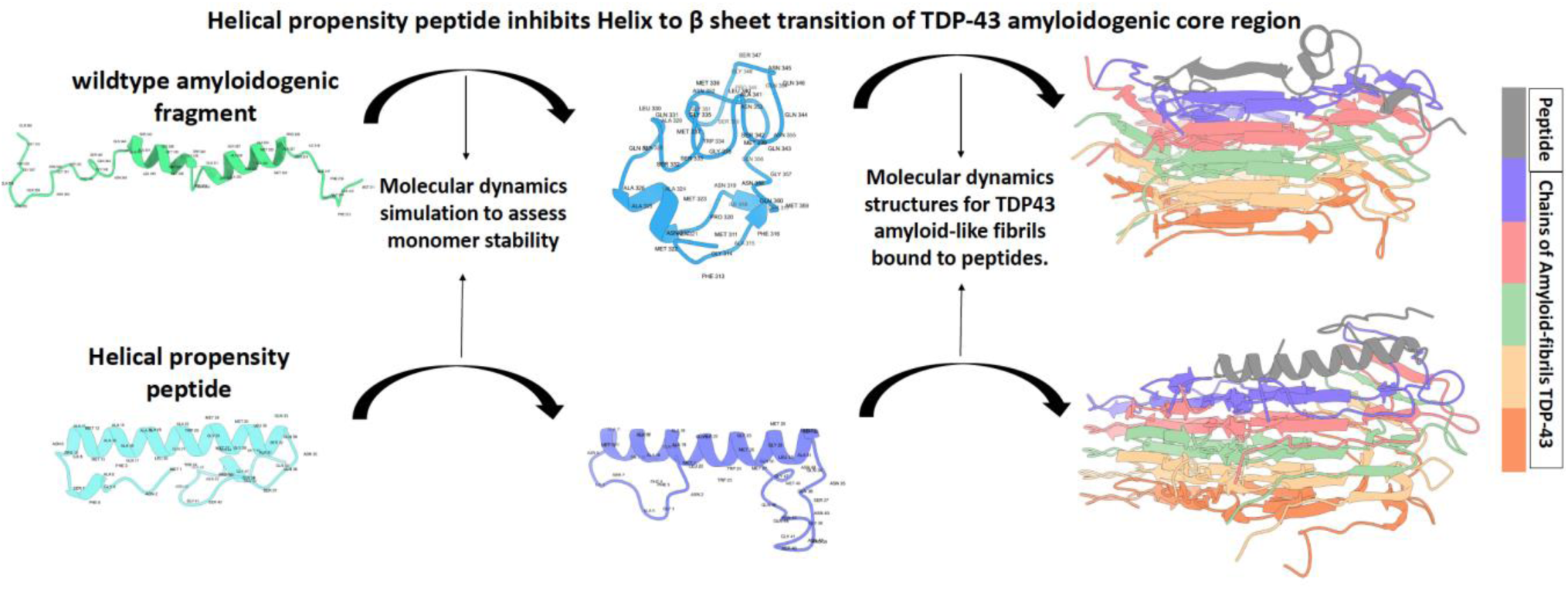

## Introduction

Amyotrophic Lateral Sclerosis (ALS) is a neurodegenerative disorder characterized by loss of motor neurons in the brain and spinal cord, leading to progressive muscle weakness, paralysis, and eventually death.^1,2^ A critical feature of ALS pathophysiology is the nuclear clearance and misfolding of essential RNA-binding proteins, resulting in the formation of insoluble aggregates.^3,4^ Nearly all (97%) ALS cases show pathologic aggregation and nuclear depletion of transactive response DNA-binding protein of 43 kDa (TDP-43) a highly conserved RNA/DNA-binding protein that plays a crucial role in RNA splicing and repression of cryptic exons, regulation of mRNA stability, and transport.^5–10^ Familial forms of ALS (5-10% of cases) have been linked to mutations in several genes, including *TARDBP* which encodes TDP-43.^11,12^ Pathological mislocalization and aggregation of TDP-43 is also a hallmark of Limbic-predominant age-related TDP-43 encephalopathy (LATE), a recently characterized form of dementia,^13^ ~50% of frontotemporal lobar degeneration (FTLD-TDP), and has been observed in a subset of Alzheimer’s disease (AD) cases.^3,14–17^ The broad relevance of TDP-43 pathology to multiple age-related neurodegenerative disorders underscores the need to elucidate molecular and/or structural alterations that trigger TDP-43 aggregation and to develop therapies targeting these pathways.

TDP-43 is a multi-domain DNA/RNA-binding protein that comprises a folded N-terminal domain (residues 1-102) important for nuclear localization, oligomerization and mediating protein-protein interactions;^18–22^ two RNA recognition motifs (RRM1, residues 104-176; and RRM2, residues 192-262) critical for binding to UG-rich RNA sequences;^23,24^ and a predominantly disordered C-terminal low complexity domain (residues 274-414).^25,26^ The C-terminal low-complexity domain (LCD) of TDP-43, also referred to as a prion-like domain, is enriched in glycine, glutamine, and asparagine residues, and is a hotspot for ALS- and frontotemporal dementia (FTD)-associated mutations.^27^ This region regulates TDP-43 solubility and serves as a crucial site for protein-protein interactions necessary for splicing, auto-regulation and other physiologic functions.^6,19,28–32^ Though predominantly disordered, the LCD also contains an α-helical domain (aa 320–340) divided into two helical segments: residues 321-330 (*AMMAAAQAAL*) and 335-343 (*GMMGMLASQ*) ***(Fig. 1A)***. Intermolecular interactions of the helical domain play an important role in driving TDP-43 liquid-liquid phase separation and assembly within physiologic ribonucleoprotein condensates in cells.^33–36^ However, under pathological conditions, the LCD is abnormally cleaved to form insoluble granulo-filamentous aggregates and significantly contributes to the process of TDP-43 aggregation.^17,25,37–41^

**Figure 1.**
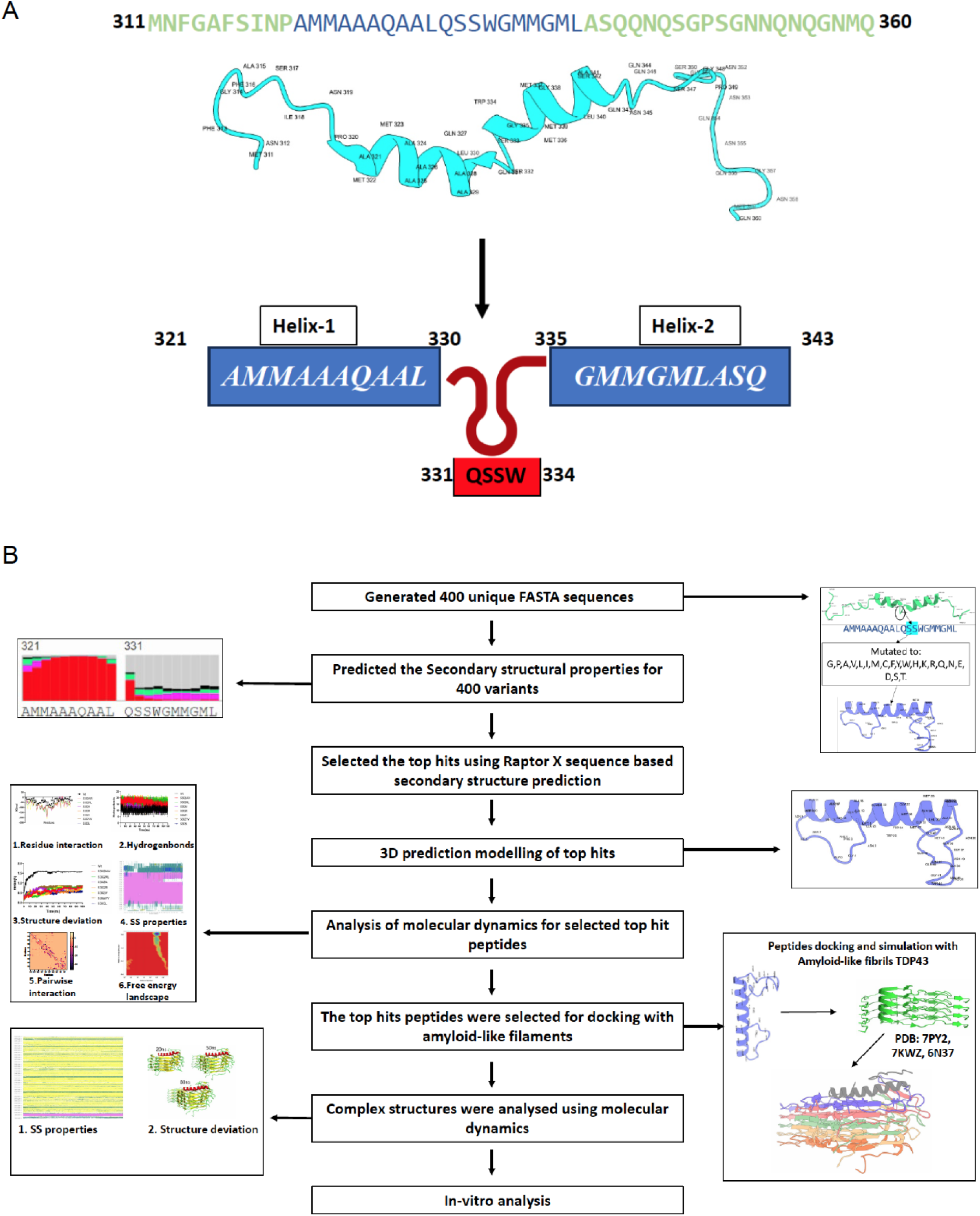
(**A**) Schematic of TDP-43’s amyloidogenic core region, highlighting crucial residues within the α-helical domain which are involved in misfolding and aggregation of TDP-43. (**B**) Schematic outline of the step-by-step approach used to develop peptides with enhanced helical propensity and analysis methods to assess peptide inhibitor stability and ability to resist beta-strand conversion.

Within the LCD, several studies have identified an amyloidogenic core region that includes the α-helical domain ***(Fig.1A)***.^42–44^ This core region (residues 311-360) is necessary and sufficient for TDP-43 aggregate formation in cells and *in vivo*, and is able to template aggregation of full-length TDP-43, suggesting a key role in regulating the structural transition of TDP-43 to amyloid-like conformations^42,45^. Indeed, cryo-electron microscopy (cryo-EM) studies of TDP-43 fibrils as well as pathological TDP-43 aggregates from the cortex of ALS/FTLD patients support the importance of the amyloidogenic core residues and α-helical domain.^46–48^ Cryo-EM of patient-derived aggregates revealed that the ordered filament core, composed of stacked TDP-43 molecules, adopts a double-spiral fold that is formed by the amyloidogenic core and flanking residues (282-360). Furthermore, the nucleus of the double-spiral fold contains several β-strands, formed by hydrophobic residues of the α-helical domain.^46^

Recent NMR studies and biophysical assays have examined how the α-helical region within the amyloidogenic core interacts with neighboring residues and changes structure during the early stages of aggregation.^42,49^ Specifically, residues 331-334 (*QSSW*) ***(Fig.1A)*** that divide the α-helical domain into two helical segments play a key role in the conversion of intramolecular helix-helix contacts (i.e., interactions between amino acid side chains) to helix-beta sheet contacts. Solid state NMR studies identified a low propensity for helical conformational chemical shift at the Ser 332 residue,^50^ and S332 in particular was implicated in many inter-residue contacts that led to fibrillar conversion of helix to beta sheet fibrils.^49^ In further support of this notion, phosphorylation of serine residues (S332, S333) also leads to helical breakage, TDP-43 misfolding and aggregation.^49^ Taken together, these data suggest the lack of helix stabilizing intramolecular contacts between S332, S333 and neighboring residues within the adjacent helical segments contribute to protein misfolding and structural transformation into amyloid-like fragments.^42,49^

Based on these findings and cryo-EM structures of TDP-43 protofilaments, in this study we used computational and rational design techniques to generate novel peptides with enhanced helical propensity that bind to TDP-43 amyloid-like filaments but resist beta sheet conversion. This strategy effectively caps the growing tips of filaments (see graphical abstract), and similar methods have been used to design peptide-based inhibitors of tau, β amyloid, and synuclein.^51^ Using the sequence of TDP-43’s amyloidogenic core (residues 311-360) as a starting point, we generated 400 novel peptides (50 residues in length) with mutations that stabilize the α-helical region. We hypothesized that peptides with mutations of S332 and/or S333 which enhance α-helix stability, would bind TDP-43 amyloid-like filaments but resist conversion to beta-sheet structure. We then employed multiple computational methodologies, including sequence-based secondary structure property prediction, three dimensional (3D)-structure modeling of peptides, docking analysis, and molecular dynamics to rank the novel peptides according to their helical secondary structure, binding efficiency with TDP-43 amyloid-like fibrils and ability to resist structural transformation ***(Fig.1B)***. Our approach takes advantage of recent advances in computing power and innovations in *in-silico* techniques to rationally design and test peptide inhibitors with therapeutic potential against TDP-43 aggregation-related disorders. This strategy may be used more broadly for *de novo* therapeutic peptide design.

## Results and Discussion

### Computational Design Pipeline

Considering the critical role of the amyloidogenic core and α-helical domain residues in initiating TDP-43 aggregation, we used an *in silico* approach to investigate whether peptide inhibitors with enhanced α-helical propensity are a potential therapeutic strategy to inhibit the growth of pathological TDP-43 filaments ***(Fig.1B)***. We retrieved the wild-type sequence of TDP-43 (residues 311-360) from the Uniprot database (Q13148) and generated a library of unique peptides each with a length of 50 residues. Given the helix destabilizing effects of serine residues 332 and 333,^50^ we focused on peptides with modifications of these polar residues. Helical secondary structure is characterized by robust intramolecular interactions; therefore, we reasoned that substitutions at S332 and/or S333 could potentially increase interactions that contribute to the stability of the helical domain. This led us to concentrate on 400 distinct mutant peptide variants with diverse secondary structural properties. RaptorX, an ultra-deep convolutional residual neural network that makes structure predictions from the primary sequence, was used to predict the secondary structure of each variant in our peptide library.^52^ Using this approach, we generated a ranked list of peptides according to their helical propensity **(Supplementary table 1)**. We found that peptides with substitutions of S332A/S333W, S332R/S333L, S332A, S332R, S332V/S333V, S332V, and S333L exhibited an enhanced helical propensity compared to the wild-type amyloidogenic core peptide sequence. These top peptide candidates were analyzed further with 3D-modeling, docking analysis, and molecular dynamics.

### Increased intramolecular interactions stabilize the helical structure in top hit peptide variants

To gain a deeper understanding of the structural characteristics of the top peptide hits, we used Rosetta comparative modeling (RosettaCM) to generate 3D-models of backbone and side-chain atom topologies.^53^ The predicted 3D structures were used to analyze the number and stability of intramolecular interactions, structural behavior, and differential energy profiles at each residue ***(Fig. 2)***. ***Fig 2A*** depicts the 3D structural coordinates of the top peptide hits, highlighting their increased helical propensity compared to the wild-type model. In order to understand the basis of enhanced helical structure of our top peptide hits, we analyzed the number of hydrogen bonds for each peptide. Notably, the helical propensity variants displayed an increased number of intramolecular hydrogen bonds compared to the wild-type peptide (***Fig. 2A)***. Overall, 22 hydrogen bonds were maintained in the wild-type helical domain, whereas nearly 32 hydrogen bonds were maintained in the helical propensity peptides helical domain (Table 1).

**Figure 2.**
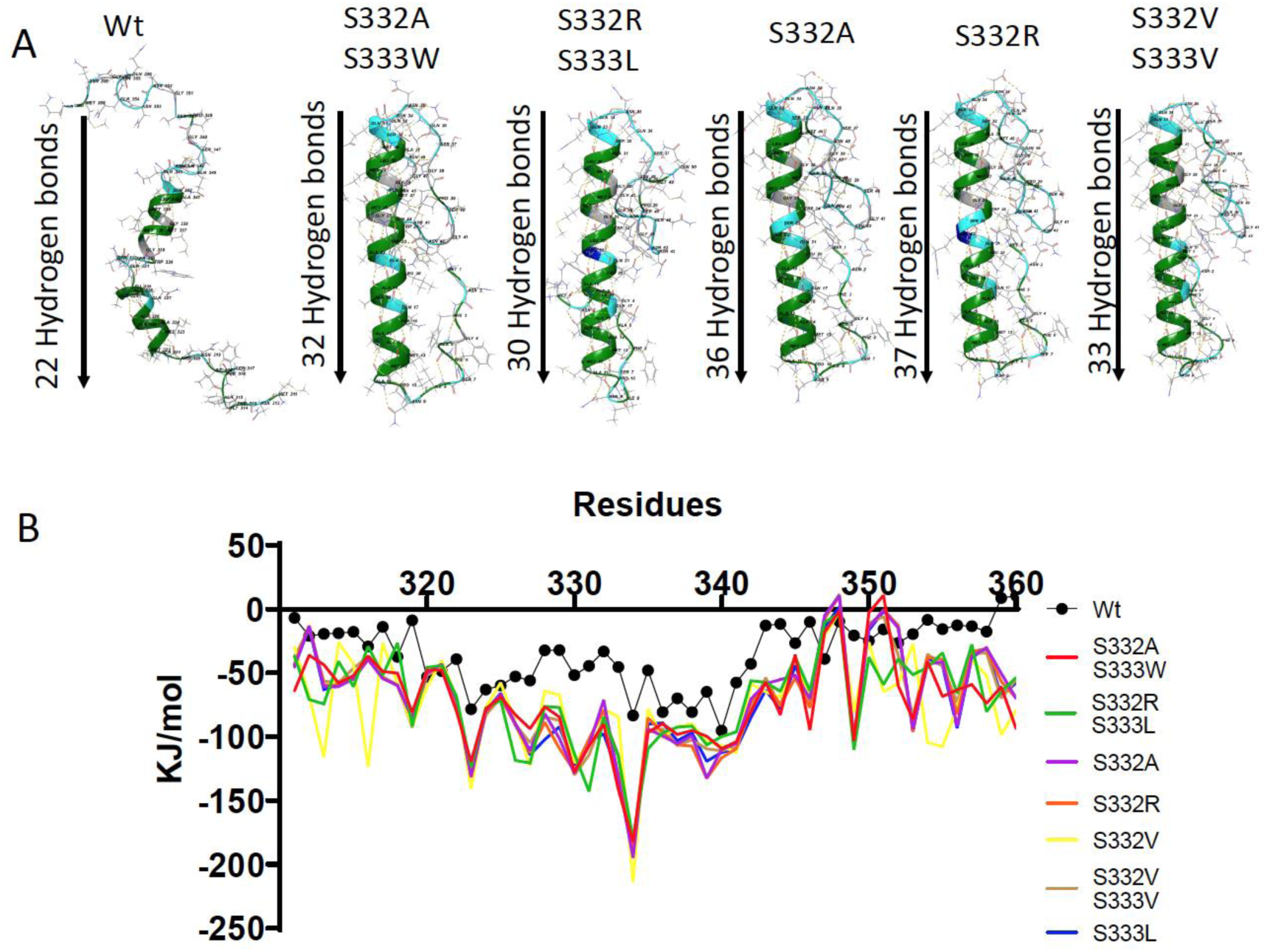
Comparative analysis of wild type (Wt) and helical propensity variant peptide structures and intramolecular interaction energies. (**A**) 3D structure representations of Wt and top hit peptides showing enhanced helical propensity. Comparison of the number of intra-molecular hydrogen bonds within the helical region underscores the structural differences between Wt and the top helical propensity peptides. (**B**) Intra-molecular interaction energy is plotted for Wt (black) and helical propensity peptides at each residue. The higher (more negative) differential energy profiles for the top hit peptides offers insight into their stability and illustrates different energetic contributions at specific residues.

**Table 1:** Number of hydrogen bonds and contributing atoms in helical domain for Wt and helical propensity variants.

| S.no | Peptides | Intramolecular interaction in the helical domain | No.of hydrogen bonds |
| --- | --- | --- | --- |
| 1 | Wt | A:MET323:H - A:PRO320:O, A:ALA324:H - A:PRO320:O, A:ALA325:H - A:ALA321:O, A:ALA326:H - A:MET322:O, A:GLN327:H - A:MET323:O, A:ALA328:H - A:ALA324:O, A:ALA328:H - A:ALA325:O, A:ALA329:H - A:ALA325:O, A:ALA329:H - A:ALA326:O, A:LEU330:H - A:ALA326:O, A:GLN331:H - A:ALA328:O, A:SER333:H - A:LEU330:O, A:TRP334:H - A:LEU330:O, A:TRP334:H - A:SER333:OG, A:GLY335:H - A:LEU330:O, A:MET336:H - A:GLN331:O, A:MET337:H -, A:TRP334:O, A:GLY338:H - A:GLY335:O, A:MET339:H - A:GLY335:O, A:LEU340:H -A:MET336:O, A:PRO320:HD2 -A:PRO320:O, A:GLY335:HA3 - A:GLN331:O | 22 |
| 2 | S332A<br>S333W | A:ALA14:H - A:PRO10:O, A:ALA15:H - A:ALA11:O<br>A:ALA16:H - A:MET12:O, A:ALA16:H - A:MET13:O<br>A:GLN17:H - A:MET13:O, A:ALA18:H - A:ALA14:O<br>A:ALA19:H - A:ALA15:O, A:LEU20:H - A:ALA16:O<br>A:LEU20:H - A:GLN17:O, A:GLN21:H - A:GLN17:O<br>A:GLN21:H - A:ALA18:O, A:ALA22:H - A:ALA18:O<br>A:ALA22:H - A:ALA19:O, A:TRP23:H - A:ALA19:O<br>A:TRP23:H - A:LEU20:O, A:TRP24:H - A:LEU20:O<br>A:GLY25:H - A:GLN21:O, A:MET26:H - A:ALA22:O<br>A:MET27:H - A:TRP23:O, A:GLY28:H - A:TRP24:O<br>A:MET29:H - A:GLY25:O, A:LEU30:H - A:MET26:O<br>A:ALA31:H - A:MET27:O, A:ALA31:H - A:GLY28:O<br>A:SER32:H - A:GLY28:O, A:GLN33:H - A:MET29:O<br>A:GLN33:H - A:LEU30:O, A:GLN34:H - A:LEU30:O<br>A:TRP23:HD1 - A:ALA19:O,A:GLY28:HA2 - A:TRP24:O, | 32 |
|  |  | A:SER32:HB2 - A:MET29:O |  |
| 3 | S332R<br>S333L | A:ALA15:H - A:ALA11:O,A:ALA16:H - A:MET12:O<br>A:ALA16:H - A:MET13:O, A:GLN17:H - A:MET13:O<br>A:ALA18:H - A:ALA14:O, A:ALA19:H - A:ALA15:O<br>A:LEU20:H - A:ALA16:O, A:GLN21:H - A:GLN17:O<br>A:GLN21:H - A:ALA18:O, A:ARG22:H - A:ALA18:O<br>A:LEU23:H - A:ALA19:O, A:TRP24:H - A:LEU20:O<br>A:GLY25:H - A:GLN21:O, A:MET26:H - A:ARG22:O<br>A:MET26:H - A:LEU23:O, A:MET27:H - A:LEU23:O<br>A:GLY28:H - A:TRP24:O, A:GLY28:H - A:GLY25:O<br>A:MET29:H - A:GLY25:O, A:MET29:H - A:MET26:O<br>A:LEU30:H - A:MET26:O, A:ALA31:H - A:MET27:O<br>A:ALA31:H - A:GLY28:O, A:SER32:H - A:GLY28:O<br>A:GLN33:H - A:MET29:O, A:GLN33:H - A:LEU30:O<br>A:GLN34:H - A:LEU30:O, A:GLN34:H - A:ALA31:O<br>A:GLY28:HA2 - A:TRP24:O, A:SER32:HB2 - A:MET29:O | 30 |
| 4 | S332A | A:ALA14:H - A:PRO10:O, A:ALA15:H - A:ALA11:O<br>A:ALA16:H - A:MET12:O, A:ALA16:H - A:MET13:O<br>A:GLN17:H - A:MET13:O, A:GLN17:H - A:ALA14:O<br>A:ALA18:H - A:ALA14:O, A:ALA19:H - A:ALA15:O<br>A:LEU20:H - A:ALA16:O, A:LEU20:H - A:GLN17:O<br>A:GLN21:H - A:GLN17:O, A:GLN21:H - A:ALA18:O<br>A:ALA22:H - A:ALA18:O, A:SER23:H - A:ALA19:O<br>A:SER23:H - A:LEU20:O, A:SER23:HG - A:ALA19:O<br>A:TRP24:H - A:LEU20:O, A:GLY25:H - A:GLN21:O<br>A:MET26:H - A:ALA22:O, A:MET26:H - A:SER23:O<br>A:MET27:H - A:SER23:O, A:GLY28:H - A:TRP24:O<br>A:GLY28:H - A:GLY25:O, A:MET29:H - A:GLY25:O<br>A:MET29:H - A:MET26:O, A:LEU30:H - A:MET26:O<br>A:ALA31:H - A:MET27:O, A:ALA31:H - A:GLY28:O<br>A:SER32:H - A:GLY28:O, A:SER32:H - A:MET29:O<br>A:SER32:HG - A:MET29:O, A:GLN33:H - A:MET29:O<br>A:GLN33:H - A:LEU30:O, A:ALA14:HA - A:GLN17:OE1<br>A:SER23:HB1 - A:ALA19:O, A:SER23:HB1 - A:LEU20:O | 36 |
| 5 | S332R | A:ALA14:H - A:PRO10:O, A:ALA15:H - A:ALA11:O<br>A:ALA16:H - A:MET12:O, A:ALA16:H - A:MET13:O<br>A:GLN17:H - A:MET13:O, A:ALA18:H - A:ALA14:O<br>A:ALA19:H - A:ALA15:O, A:LEU20:H - A:ALA16:O<br>A:LEU20:H - A:GLN17:O, A:GLN21:H - A:GLN17:O<br>A:GLN21:H - A:ALA18:O, A:ARG22:H - A:ALA18:O<br>A:SER23:H - A:ALA19:O, A:SER23:H - A:LEU20:O<br>A:SER23:HG - A:ALA19:O, A:TRP24:H - A:LEU20:O<br>A:GLY25:H - A:GLN21:O, A:MET26:H - A:ARG22:O<br>A:MET26:H - A:SER23:O, A:MET27:H - A:SER23:O<br>A:GLY28:H - A:TRP24:O, A:GLY28:H - A:GLY25:O<br>A:MET29:H - A:GLY25:O, A:MET29:H - A:MET26:O<br>A:LEU30:H - A:MET26:O, A:ALA31:H - A:MET27:O<br>A:ALA31:H - A:GLY28:O, A:SER32:H - A:GLY28:O<br>A:SER32:H - A:MET29:O, A:SER32:HG - A:MET29:O | 37 |
|  |  | A:GLN33:H - A:MET29:O, A:GLN33:H - A:LEU30:O<br>A:GLN34:H - A:LEU30:O, A:GLN34:H - A:ALA31:O<br>A:ALA14:HA - A:GLN17:OE1, A:SER23:HB1 - A:ALA19:O<br>A:SER23:HB1 - A:LEU20:O |  |
| 6 | S332V<br>S333V | A:ALA14:H - A:PRO10:O, A:ALA15:H - A:ALA11:O<br>A:ALA16:H - A:MET12:O, A:ALA16:H - A:MET13:O<br>A:GLN17:H - A:MET13:O, A:ALA18:H - A:ALA14:O<br>A:ALA19:H - A:ALA15:O, A:LEU20:H - A:ALA16:O<br>A:LEU20:H - A:GLN17:O, A:GLN21:H - A:GLN17:O<br>A:GLN21:H - A:ALA18:O, A:VAL22:H - A:ALA18:O<br>A:VAL22:H - A:ALA19:O, A:VAL23:H - A:ALA19:O<br>A:VAL23:H - A:LEU20:O, A:TRP24:H - A:LEU20:O<br>A:GLY25:H - A:GLN21:O, A:GLY25:H - A:VAL22:O<br>A:MET26:H - A:VAL22:O, A:MET27:H - A:VAL23:O<br>A:GLY28:H - A:TRP24:O, A:GLY28:H - A:GLY25:O<br>A:MET29:H - A:GLY25:O, A:MET29:H - A:MET26:O<br>A:LEU30:H - A:MET26:O, A:ALA31:H - A:MET27:O<br>A:ALA31:H - A:GLY28:O, A:SER32:H - A:GLY28:O<br>A:GLN33:H - A:MET29:O, A:GLN33:H - A:LEU30:O<br>A:GLN34:H - A:LEU30:O, A:GLN34:H - A:ALA31:O<br>A:SER32:HB2 - A:MET29:O | 33 |
| 7 | S333L | A:ALA14:H - A:PRO10:O, A:ALA15:H - A:ALA11:O<br>A:ALA16:H - A:MET12:O, A:ALA16:H - A:MET13:O<br>A:GLN17:H - A:MET13:O, A:ALA18:H - A:ALA14:O<br>A:ALA19:H - A:ALA15:O, A:ALA19:H - A:ALA16:O<br>A:LEU20:H - A:ALA16:O, A:GLN21:H - A:GLN17:O<br>A:SER22:H - A:ALA18:O, A:SER22:H - A:ALA19:O<br>A:SER22:HG - A:ALA18:O, A:LEU23:H - A:ALA19:O<br>A:LEU23:H - A:LEU20:O, A:TRP24:H - A:LEU20:O<br>A:GLY25:H - A:GLN21:O, A:MET26:H - A:SER22:O<br>A:MET27:H - A:LEU23:O, A:GLY28:H - A:TRP24:O<br>A:MET29:H - A:GLY25:O, A:MET29:H - A:MET26:O<br>A:LEU30:H - A:MET26:O, A:ALA31:H - A:MET27:O<br>A:ALA31:H - A:GLY28:O, A:SER32:H - A:GLY28:O<br>A:SER32:H - A:MET29:O, A:GLN33:H - A:MET29:O<br>A:GLN33:H - A:LEU30:O, A:GLN34:H - A:LEU30:O<br>A:GLN34:H - A:ALA31:O, A:ALA14:HA - A:GLN17:OE1<br>A:SER22:HB1 - A:ALA18:O, A:SER22:HB1 - A:ALA19:O<br>A:GLY28:HA2 - A:TRP24:O, A:SER32:HB2 - A:MET29:O | 36 |
| 8 | S332V | A:ALA14:H - A:PRO10:O, A:ALA15:H - A:ALA11:O<br>A:ALA16:H - A:MET12:O, A:GLN17:H - A:MET13:O<br>A:GLN17:HE21 - A:GLN21:OE1, A:ALA18:H - A:ALA14:O<br>A:ALA19:H - A:ALA15:O, A:ALA19:H - A:ALA16:O<br>A:LEU20:H - A:ALA16:O, A:LEU20:H - A:GLN17:O<br>A:GLN21:H - A:GLN17:O, A:VAL22:H - A:ALA18:O<br>A:VAL22:H - A:ALA19:O, A:SER23:H - A:ALA19:O<br>A:SER23:H - A:LEU20:O, A:TRP24:H - A:LEU20:O<br>A:GLY25:H - A:GLN21:O, A:MET26:H - A:VAL22:O<br>A:MET26:H - A:SER23:O, A:MET27:H - A:SER23:O<br>A:MET27:H - A:TRP24:O, A:GLY28:H - A:TRP24:O<br>A:MET29:H - A:GLY25:O, A:LEU30:H - A:MET26:O<br>A:ALA31:H - A:MET27:O, A:ALA31:H - A:GLY28:O<br>A:SER32:H - A:GLY28:O, A:GLN33:H - A:MET29:O<br>A:ALA18:HA - A:GLN17:OE1, A:SER23:HB1 - A:LEU20:O<br>A:SER32:HB1 - A:GLY28:O, A:SER32:HB2 - A:MET29:O | 32 |

Beyond the analysis of hydrogen bonds, our investigation sought to comprehensively assess the various energy interaction types within these peptides by utilizing an interaction energy matrix.^54^ This encompassed a comprehensive prediction of intramolecular interaction energies, including all categories of covalent, polar, and non-polar interactions. This analysis demonstrated distinct values within the helical domain (residues 320-340) for the helical propensity variants and the wild-type peptide. Specifically, the helical propensity variants exhibit an average energy of −96.119 kJ/mol, while the wild-type counterpart displays an average energy of −57.8 kJ/mol ***(Fig. 2B)***.These findings suggest increased intramolecular energy observed in the helical propensity variants plays an important role in maintaining the stability of the helical structure. Upon closer examination, we observed that the substitutions at S332 and/or S333 residues contributed a marked increase in the intramolecular interaction energy. Specifically, mutations to S332A/S332W yielded an intramolecular interaction energy of −90.14 kj/mol, whereas the wild-type model exhibited less intramolecular interactions at S332 and S333 (−33.2 kj/mol) ***(Fig. 2B)***. These data indicate that alterations in the serine residues significantly influence the intramolecular interaction characteristics in the helical region (residues 320-340). These observations also suggest the mutated serine residues in our top peptide hits demonstrate a more favorable intramolecular interaction energy profile with neighboring residues, thus contributing robust interactions that stabilize their helical secondary structure.

### Stability of Helical Propensity peptide inhibitors through Molecular Dynamics

Comprehensive molecular dynamics (MD) simulations were performed to further validate the folding and conformational stability of top hit peptides and gain insight on their dynamic behavior and interplay among constituent residues, compared to the wild-type peptide. We conducted MD simulations *by using GROMACS (version 2022)*. For the initial equilibrium system, we employed Optimized Potentials for Liquid Simulations (OPLS) force field in conjunction with the Simple Point-Charge (SPCE) water model.^55^ We analyzed the wild-type and top helical propensity monomer structures for up to 100 ns to understand the time evolution of their structural characteristics ***(Fig. 3)*** and free energy properties ***(Fig. 4)***. Root mean square deviation (RMSD) analysis was employed to assess the stability and conformational changes of the proteins. Generally, the lowest RMSD values indicate a more stable conformation, while higher or fluctuating RMSD values suggest structural changes. Notably, the helical propensity variants exhibited lower RMSD values compared to the wild-type model throughout the simulation period (***Fig.3A***), suggesting their enhanced stability. Hydrogen bonds are an important source of intramolecular interactions that maintain stable helical structure. Therefore, we also examined the total number of hydrogen bonds in the helical domain (320-340) for both the wild-type and helical propensity variants. The wild-type model exhibited an average of 10 to 15 hydrogen bonds, whereas the helical propensity variants maintained 15 to 20 hydrogen bonds over the simulation period ***(Fig 3B)***. Furthermore, we determined the secondary structure properties for the helical propensity peptides throughout the simulation period. Secondary structural analysis of the wild-type model revealed an unstable helical segment spanning residues 327 to 340, that transitioned into loop structures ***(Fig 3C)***. In contrast, the helical propensity variants, such as S332A/S333W, S332R/S333L, and S333L, maintained stable secondary structures characterized by well-defined helical segments throughout the simulation. Combining all of these results, MD data indicate that substitutions of serine residues in our top peptide hits substantially enhance helical conformational stability, in part by contributing robust intramolecular hydrogen bonds that stabilize their helical secondary structure.

**Figure 3.**
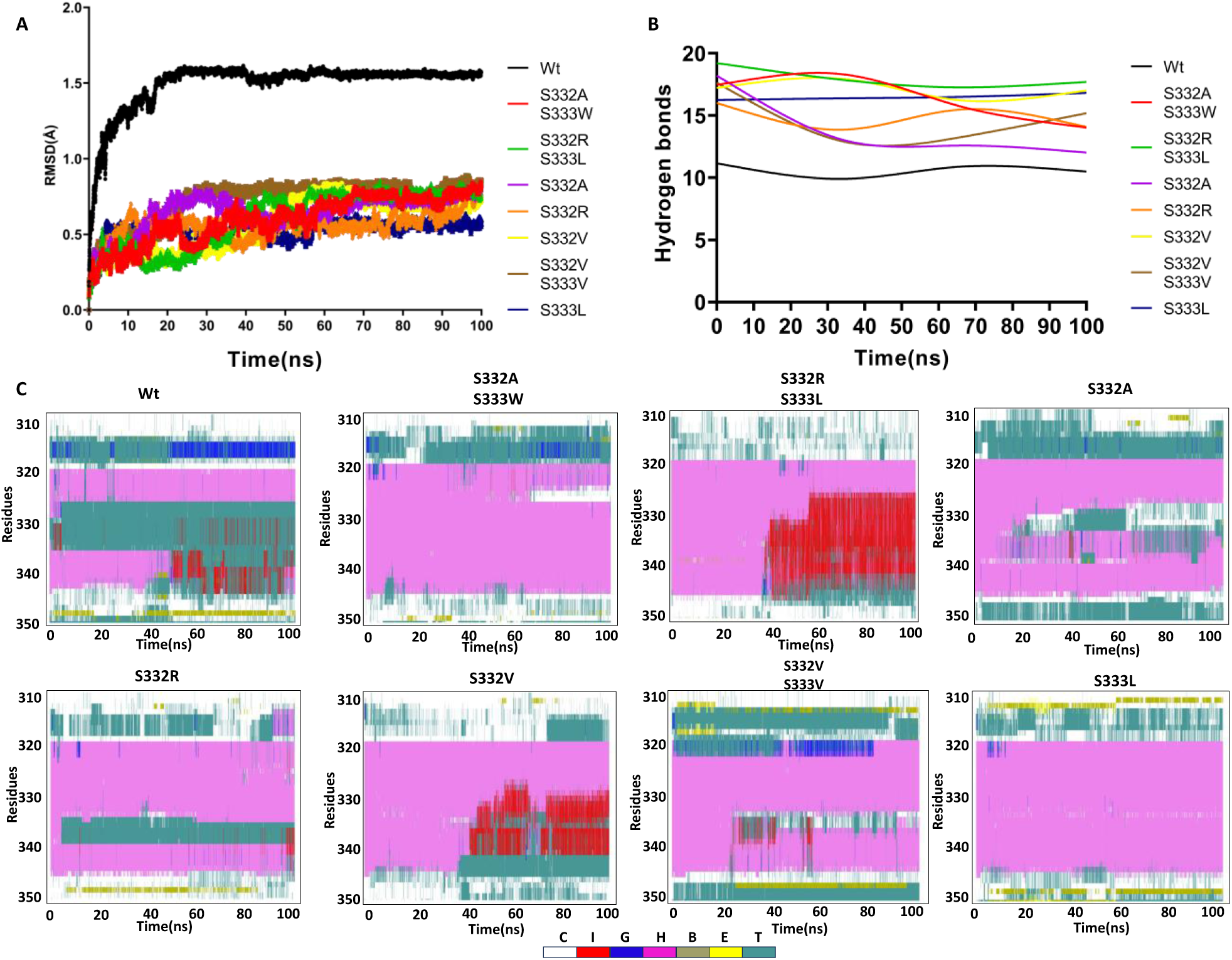
Molecular Dynamics simulations were used to assess the conformational stability of helical propensity and wild-type peptides. (**A**) The RMSD values were calculated to measure stability and conformational changes of helical propensity peptides and the wild-type model over a 100ns simulation period. The plot shows the relative stability of the helical propensity variants compared to wild-type peptide. (**B**) The total number of hydrogen bonds formed within the helical domain of each peptide was monitored throughout the 100ns simulation period. This spline plot illustrates the variation in the hydrogen bonds for the helical propensity variants and the wild-type model, providing insights into the strength and stability of the intra-molecular interactions. (**C**) Residue-level Secondary Structure Analysis for each peptide over the simulation. Secondary structural properties (C-coil, I-5turn helix, G-3turn helix, H-4turn helix, B-Beta bridge, E-Beta sheet, T-turn) were analyzed for each residue in the helical propensity variants and the wild-type model over the course of the 100 ns simulation. This plot illustrates stability of helical conformations in our top hit peptides, relative to the wild-type amyloidogenic peptide.

**Figure 4.**
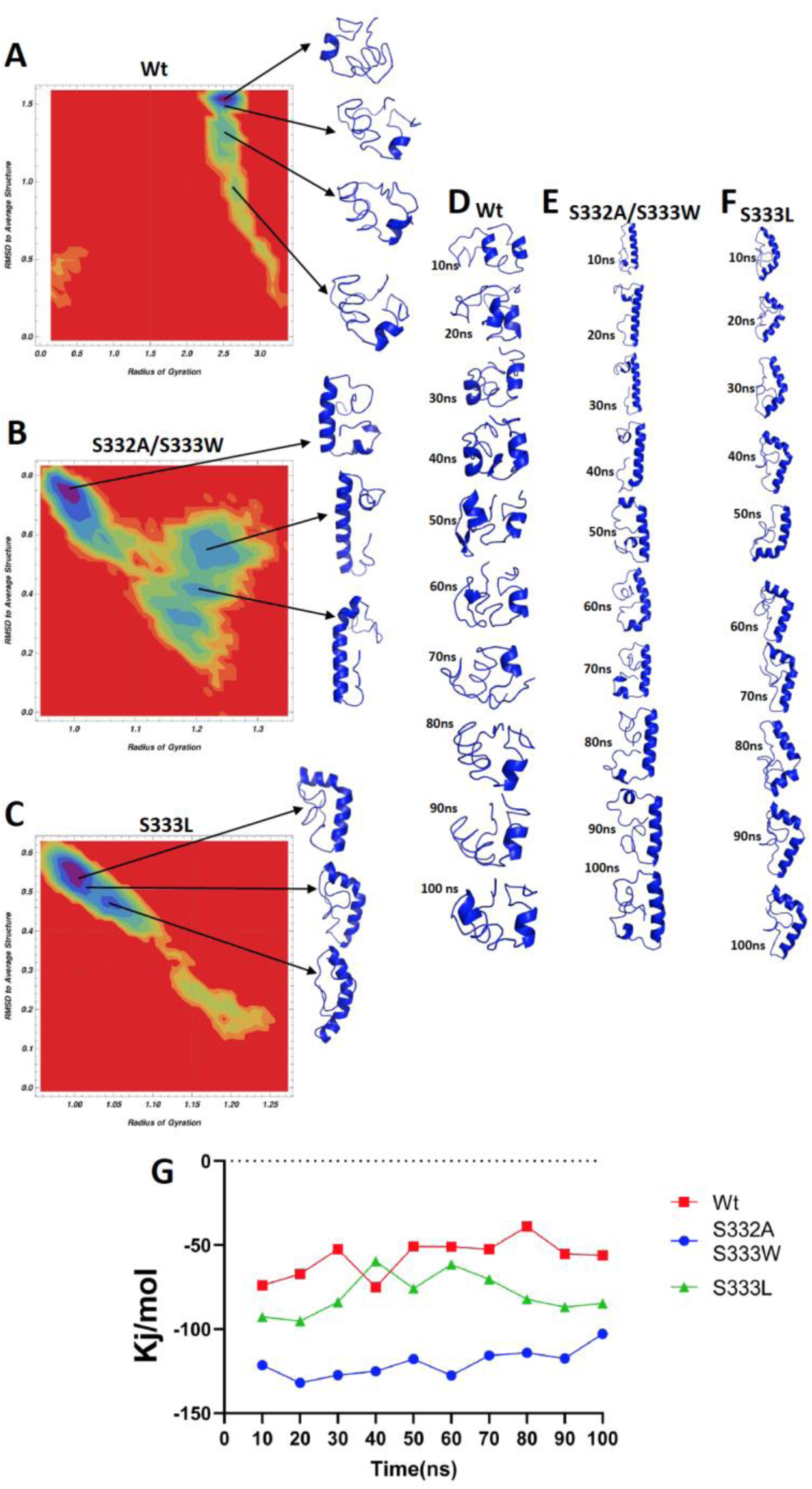
Comparative Gibbs free energy analysis from molecular dynamics simulations of wild-type (Wt) and helical propensity peptides. (**A-C**) 3D-graphical representation of calculated Gibbs free energy for (**A**) Wt, (**B**) S332A/S333W, and (**C**) and S333L peptides, providing insights into the thermodynamic stability of these structures (blue color ranges indicate a favorable, high Gibbs free energy and red color ranges indicate low Gibbs free energy). (**D-F**) Selected snapshots from molecular dynamics simulations at various timepoints for (**D**) Wt, (**E**) S332A/S333W, and (**F**) S333L peptides, illustrating the dynamic behavior and conformational changes over time. (**G**) The cumulative contribution of intramolecular interaction energies between residues at 332 and 333 position and flanking neighboring residues (A329, L330, Q331, W334, G335, and M336) observed over the course of 100ns MD. Analysis performed for wild-type (S332, S333) model and peptide variants with mutated residues S332A/S33W and S333L.

Next, we predicted thermodynamic stability of the wild-type and top two helical propensity peptides over the simulation period. From the MD simulations, Gibbs free energy heatmaps were generated ***(Fig. 4A-C)***. We found distinct clusters representing different conformational states of the peptides, with cooler blue colors indicating relatively stable states. The helical propensity variants, S332A/S333W and S333L, exhibited stable conformations throughout the simulation, as indicated by the favorable Gibbs free energy values within their respective clusters ***(Fig. 4B, 4C)***. In contrast, the wild-type model showed structural changes at 1.5Å RMSD, at which point it reached an unstable state with reduced free energy ***(Fig. 4A, 4D)***. This instability was mainly attributed to the presence of polar residues, such as S332 and S333, in the helical domain, resulting in fewer hydrogen bonds and unstable intramolecular interaction energy. Conversely, the helical propensity variants maintained stable conformations with stable intramolecular interaction energies ***(Fig. 2B; Supplementary Fig. 1)***.

### The Role of Serine 332,333 Residues in TDP-43 Fibrillization: Insights into the Importance of Neighbor Interactions in the Helical Domain

Previous investigations have pinpointed interresidue interactions, particularly interactions involving S332 and W334, that contribute to structural conformational changes of TDP-43’s amyloidogenic core and subsequent fibril formation.^49,56^ It has been suggested by Zhuo et al that the lack of intramolecular interactions between S332 and neighboring residues has a potential impact on fibril initiation.^49^ SSNMR with proton detection also revealed a close interaction between the indole Nε1–Hε1 of W334 and the side-chain carbonyl of Q343. Given the fundamental requirement of 3 to 4 residues for sustaining a single turn within the helical structural motif, we investigated how substitutions of S332 and/or S333 in our top hit peptides impact the cumulative intramolecular interaction energies between these residues and three flanking residues in the helical domain (A329, L330, Q331, W334, G335, and M336). Over the course of a 100ns simulation period, it became evident that the wild-type model exhibited a less favorable cumulative intramolecular interaction energy profile compared to S332A/S333W and S333L peptides (***Fig 4G***). These findings support the contribution of robust helix-stabilizing intramolecular interactions seen with S332A/S333W and S333L substitutions in our top hit helical propensity peptides.

We went on to examine pairwise interactions between the residues comprising TDP-43’s amyloidogenic core region ***(Fig. 5)***. In the wild-type peptide, we observed that S332 and W334 residues from the helical domain engage in interactions with residues outside of the helical domain, particularly residues G357 and N358 (***Fig. 5A***). This observation underscores the lack of helix-stabilizing interactions between S332 and neighboring residues within the helical domain. These findings are also consistent with prior studies which show that S332 and W334 participate instead in helix-destabilizing interactions that initiate helix to beta sheet conversion.^42,49^ In contrast pairwise interactions for the S332A/S333W helical propensity peptide are most robust within the helical regions (***Fig. 5B***).

**Figure 5.**
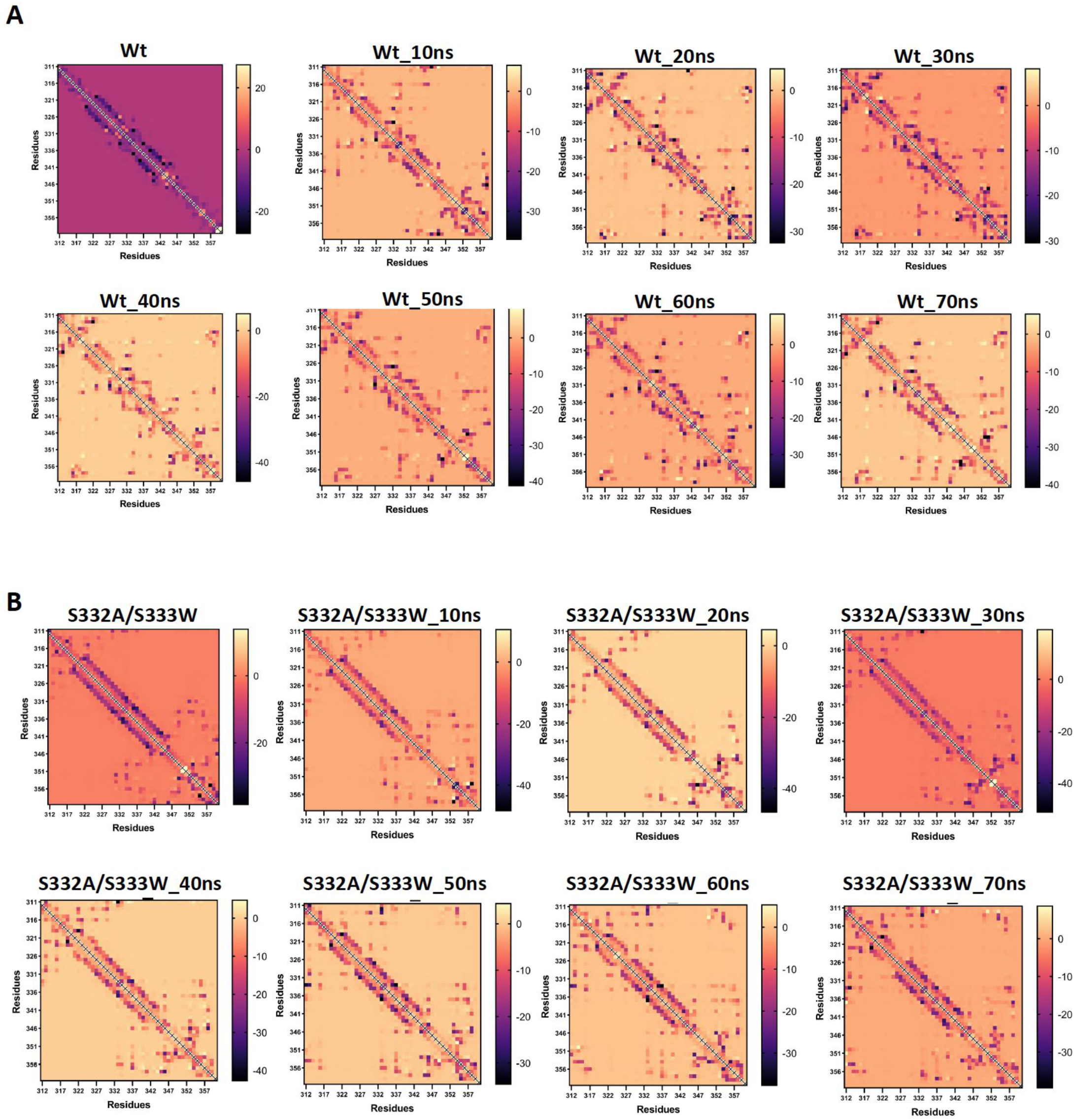
Heatmap for Predicted Pairwise Interactions of residues 311-360 across Molecular Dynamics Snapshots for (**A**) wild-type and (**B**) S332A/S333W mutant. The color gradient indicates variation in intramolecular interaction energies (kj/mol).

Furthermore, we analyzed the probability of helical secondary structure throughout the simulation period for wild-type and our helical propensity peptides. We found that the wild-type model has reduced helical probability for serine neighbor residues such as 331 to 335 ***(Supplementary Fig. 2A)***. Therefore, additional evidence from this probability data has been suggested that serine residues in the helical domain is a major residue to breaking the helix into two segments or initiating misfolding. Whereas, the S332A/S333W, S332R, S332V, S332V/S333V helical propensity peptides have higher probability compared to wild-type and other helical propensity peptides ***(Supplementary Fig. 2A)***. In addition to that, residues wise helical probability for wildtype and helical propensity peptides has shown that, the break regions has increase helical probability (***Supplementary Fig. 2B-G***). Based on these findings, the helical propensity variants were selected for further analysis, including docking studies and complex simulations with amyloid-like fibrils.

### Efficiency of Helical Propensity Peptide Binding with TDP-43 Amyloid-like Structures

The binding efficacy of diverse peptides bound to TDP-43 amyloid-like fibrils was evaluated using three distinct amyloid-like filament crystal structures of TDP-43. The first structure was derived from patient-aggregated TDP-43, located in the frontal and motor cortices of a patient diagnosed with ALS/FTLD.^46^ The additional two structures were elucidated through crystallographic analysis.^47,48^ Each of these structures displays unique beta sheet conformations. The patient-derived structure 7PY2 contains almost ten beta sheets, with the longest one extending from serine 332 to leucine 340. Meanwhile, structures 7KWZ and 6N37 exhibit ten and three beta sheets, respectively, each adopting distinct beta sheet conformations. Interestingly, the helical domain within these structures presents a lengthy beta sheet conformation. As indicated by prior studies,^42,49^ the transition step of the helical domain serves as the initial phase of aggregate formation. When taking into account these previous findings and the patient-derived and crystal-solved structures, it suggests that helical domains participate in forming the most elongated, stable beta sheets within these three structures. Therefore, we predicted that when these reference TDP-43 fibrillar structures are complexed with our top hit helical propensity peptides, we expect that our peptide inhibitors will bind amyloid-like fibrils but resist structural transformation, compared to the wild-type peptide., and furthermore, it should prevent monomer addition of TDP-43 CTD fragments to amyloid-like fibrils,

As mentioned in previous findings, the c-terminal TDP-43 always needs its interacting partners (additional C-terminal fragment) to form the fibrillation.^31^ As a result, in our previous simulation results only with monomers, we did not observe any stable beta sheet formation. Therefore, we tested this helical propensity peptide and the wildtype model with a bound stage of several crystal structures of the C-terminal TDP-43 fragment. The amyloid-like fibrils and peptides were docked using HDock and we used three different crystal structures for the docking such as 7KWZ, 7PY2 and 6N37 ***(Supplementary Fig. 3B)***. The docking analysis reveals that the helical propensity variants and wildtype fragments achieved more similar scores (**Table 2**) and the S332A/S333W helical propensity peptide is bound at the helical domain in the Amyloid like fibrils structure **(*Supplementary Fig. 3A)***. Which means, it has favorable binding affinity with amyloid-like fibrils as a TDP-43 C-terminal fragment mimetic. Overall, the binding affinity score varies between −685.51 to −759.83. To further investigate stability and structural changes between the complexes, we selected top poses with favorable interactions for the molecular dynamics simulation analysis.

**Table 2:** Docking scores of higher helical propensity variants with Amyloid like fibrils of TDP-43.

| Variants | 7KWZ (Li et al) | 7PY2 (Arseni et al) | 6N37 (Eisenberg, D.S) |
| --- | --- | --- | --- |
| S332A/S333W | -685.51 | -599.62 | -670.91 |
| S332R/S333L | -709.39 | -638.95 | -653.31 |
| S332V/S333V | -717.35 | -662.35 | -660.23 |
| S332V | -689.09 | -587.00 | -629.61 |
| S333L | -759.83 | -701.79 | -666.20 |
| S332A | -725.48 | -629.92 | -697.64 |
| S332R | -712.70 | -647.64 | -720.48 |

### Helical propensity peptides maintain helical structure and resist beta-sheet transformation when bound to TDP-43 fibrils

As a next step in testing our top peptide inhibitors, we were interested in assessing the structural stability of helical propensity peptides after binding to TDP-43 amyloid-like fibrils.. From this analysis, we aimed to find the probability of helix to beta sheet transformation of peptide monomers and after binding with amyloid-like fibrils. To elucidate the molecular interaction between the complexes, we used top poses from the docking analysis for molecular dynamics simulation. The simulation results indicate that the peptide/fibril complex formed a stable structure and maintained favorable interactions with amyloid-like fibrils. Interestingly, in the monomer state, the wild-type peptide had a more than 40% percentage reduction in helical probability compared to the S332A/S333W helical propensity peptide ***(Fig 6A, 6B)***. As we mentioned previously, we did not find any significant difference in the beta sheet probability at the monomer state ***(Fig 6C, 6D)***. Consistent with previous studies of the known amyloidogenic core, the wild-type model showed a significant increase in beta sheet probabilities and a corresponding reduction in helical probability after binding to amyloid-like fibrils ***(Fig 6E-H)***. Residue-wise analysis showed that S332 and neighbor residues, such as W334, had a marked increase in beta sheet probability and decreased helical probability ***(Fig 6E, 6G)***. On the other hand, the helical propensity peptide (S332A/S333W) maintains stable interaction with neighbor residues, and we found very low deviated values for beta sheet probability in both monomer and complex states ***(Fig 6D, 6H)***. To gain a structural view, snapshots of different time points from the molecular dynamics were examined for complexes of the S332A/S333W peptide with 7KWZ and WT_7KWZ complexes, indicating the wild-type bound model with amyloid like fibrils, no longer able to maintain the helical conformations with amyloid like fibrils, with most of the residues from the helical domain forming a loops or beta sheets ***(Supp 4a, 4c)***. Furthermore, the helical propensity peptides maintained the helical conformation throughout the simulation period ***(Supp 4b, 4c)***. We also observed some conformational changes in the helical propensity peptides during the simulation, suggesting their flexibility and potential for adopting different conformations.

**Figure 6:**
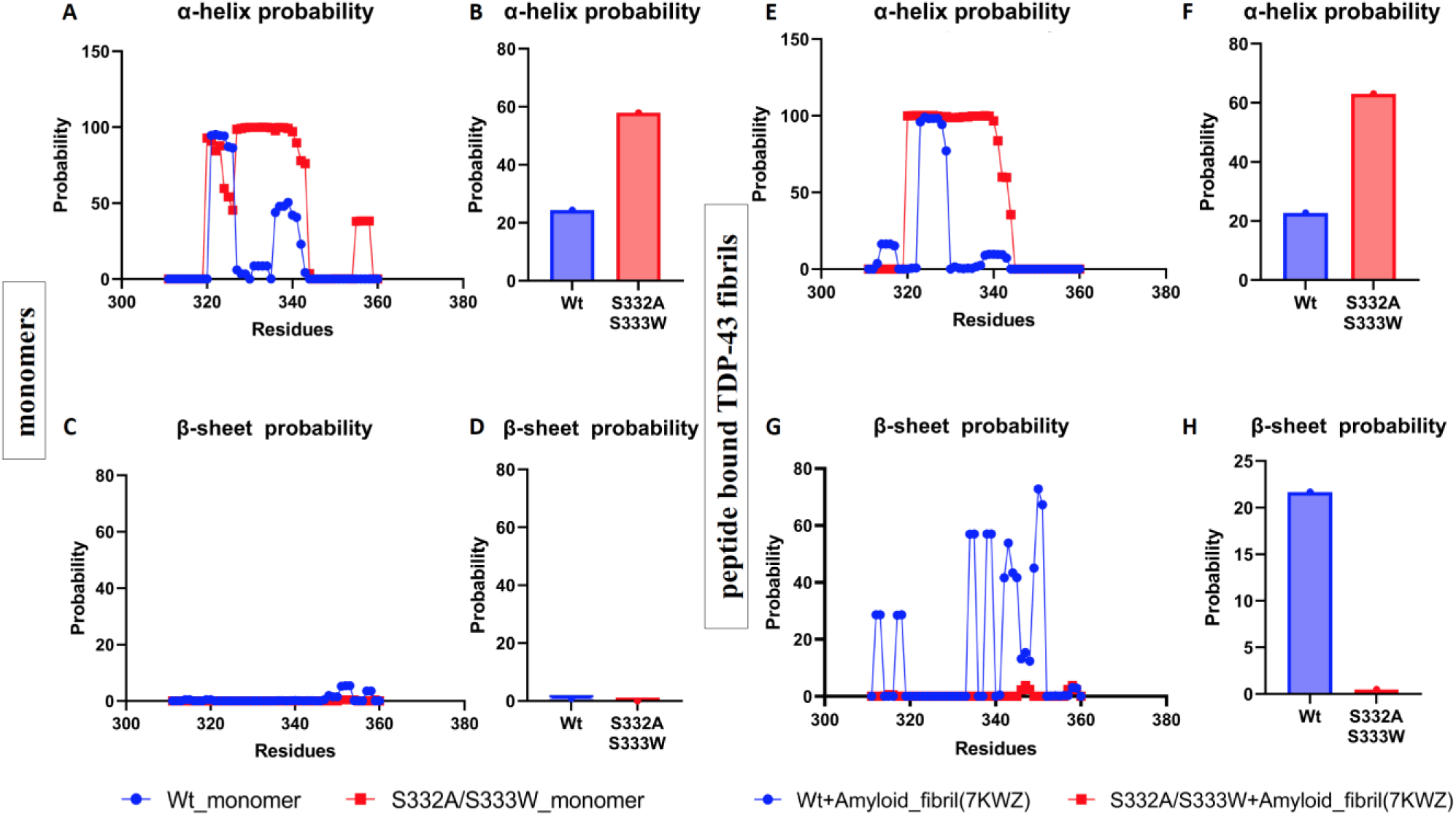
Predicted residue-wise and overall helical or beta-sheet probability for the wild-type peptide and the S332A/S333W helical probability peptide in two different states: (**A-D**) peptide monomer state and (**B-H**) complex state (peptide bound to TDP-43 amyloid-like fibrils). (**A, E**) Residue-wise predicted helical probability and (**C, G**) beta sheet probability in the monomer and complex state for the wild-type and S332A/S333W peptides. (**B, F**) Overall helical probability for the monomer state (B) and the complex state (F). (**D,H**) Overall beta sheet probability for the monomer state (D) and the complex state (H). These figures collectively provide a comprehensive analysis of the helical and beta sheet probabilities for the wild-type and helical propensity peptides in both the monomer and complex states.

### Promising Capping of TDP-43 Amyloid-Like Fibrils by Helical Propensity Peptides

Finally, we tested the thermodynamic stability of complexes of either wild-type ***(Fig. 7A,B)*** or helical propensity peptide bound to TDP-43 amyloid-like fibrils ***(Fig. 7C,D)***. To perform this analysis, we predicted the free energy landscape along with timeframe on the x-axis and helical or beta sheet probabilities on the y axis. We sought to understand whether helical propensity peptides bound to TDP-43 fibrils are more thermodynamically stable than the wild-type peptide/fibril complex. Our data suggests that the S332A/S333W peptide shows high helical probability in the complex state and was stable throughout the simulation period. For example, we found similar levels of helical probability (44-48%), was maintained from 0 to 25 and 30 to 50 ns with more free energy clusters (dark blue) ***(Fig. 7C)***. We did not detect any favorable free energy clusters for beta sheet probability ***(Fig. 7D)***. As we expected, the wild-type model, when bound to TDP-43 fibrils, has reduced helical propensity and increased beta sheet probability in high free energy clusters. Approximately 10% helical probability and more than 25% beta sheet probability structures had dark blue high free energy clusters (***Fig. 7A, B***). Overall, our data suggest that the S332A/S333W helical propensity peptide has strong binding efficiency with amyloid like fibrils, forming a thermodynamically stable complex structure, while still maintaining stable helical propensity. Thus, this rationally designed helical propensity peptide displays promising features, as it can bind and cap TDP-43 amyloid like fibrils, acts as a mimetic of TDP-43’s amyloidogenic core region, and but resists beta sheet transformation. Overall, our study provides insights into the binding mechanism and stability of TDP-43 C-terminal fragment peptide mimetics, which could be useful for developing new therapies combating TDP-43 proteinopathies.

**Figure 7:**
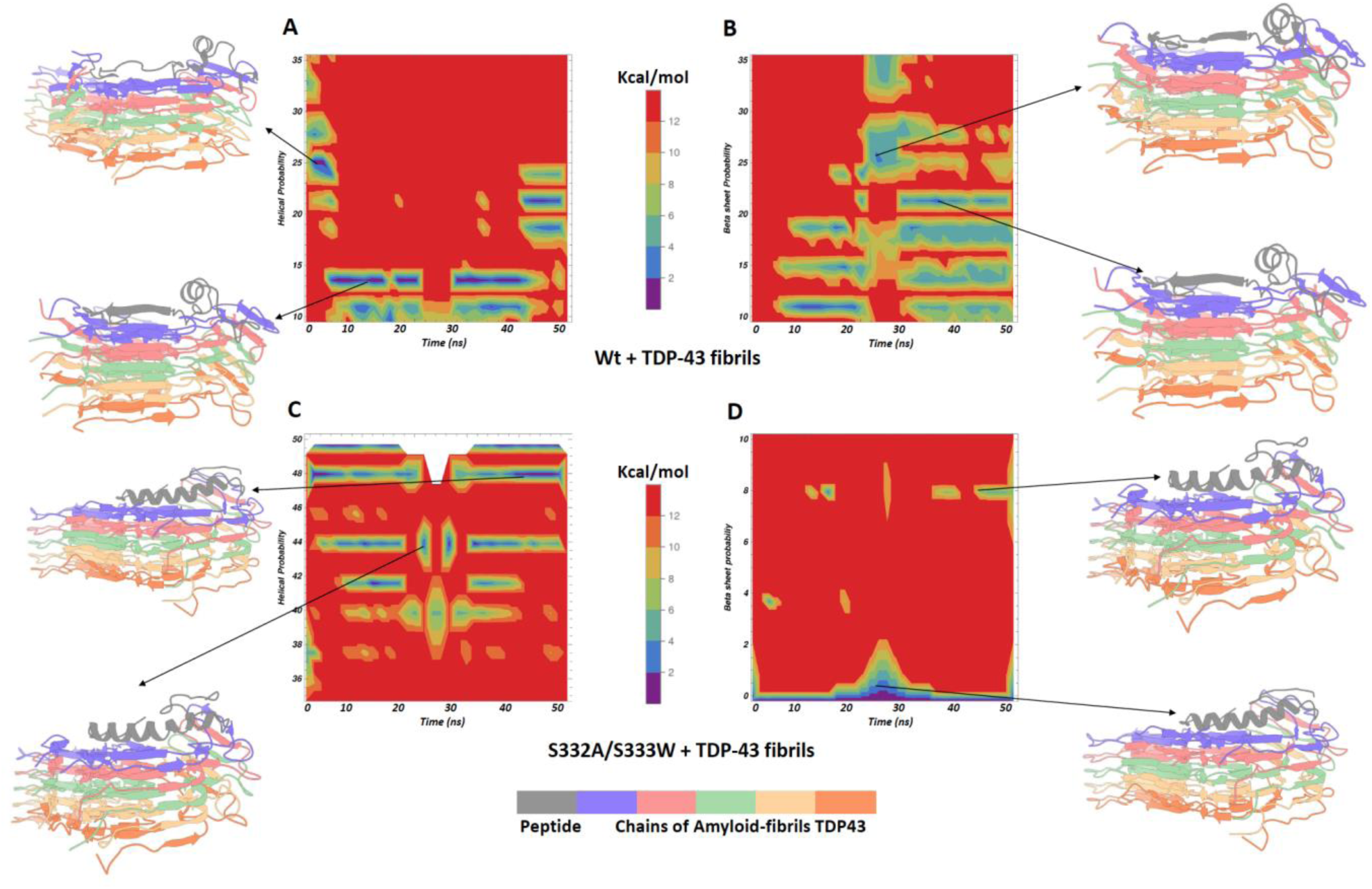
Predicted free energy landscape of (**A,B**) wild-type or (**C,D**) S332A/S333W peptide-bound amyloid-like fibrils, with time(ns) plotted along the x-axis and the helical or beta sheet probabilities represented on the y-axis. Each chain of the amyloid-like fibrils is color-coded with distinct colors, while the peptides are highlighted in gray. The free energy values are depicted using a rainbow color scheme, where favorable high-energy regions are represented in blue or violet, and unfavorable lower-energy regions are shown in orange or red. This landscape provides insights into the conformational stability and dynamics of the peptides within the amyloid-like fibril structure, showcasing conversion from helical to beta sheet conformations and their associated free energy variations.

## Conclusion

By utilizing advanced in-silico techniques we uncover a new direction for the development of therapeutic strategies against TDP-43 aggregation-related disorders, such as ALS and FTD. Our work specifically targets the destabilization of the α-helical domain in the C-terminal region of TDP-43, a process that is critical to the formation of pathological amyloid-like filaments. These helical domains act as crucial sentinels, effectively safeguarding against the formation of amyloid-like fibrils. Simultaneously, it is essential to explore the physiological functions of helical domains, ensuring that mutations within this domain do not disrupt TDP-43 transport in axons, hinder essential protein-protein interactions, impede splicing, or other vital functions. Therefore, these peptides require further evaluation for specificity and target engagement *in vitro* and in cellular assays. The peptides identified in our computational analysis, distinguished by their enhanced stability and binding affinity, are currently the subject of further validation through *in vitro* and *in vivo* experiments. These subsequent investigations are essential steps in confirming the therapeutic potential of these peptides and moving closer to the development of novel therapeutic strategies for TDP-43 aggregation-related disorders. The potential of *in silico* methodologies to accelerate the discovery of therapeutic leads, represents a significant contribution to the field of drug discovery for neurodegenerative diseases. Our findings underscore the prospect of targeting the α-helical domain in TDP-43 to prevent the formation of pathological filaments, a strategy that could potentially revolutionize the treatment of conditions such as ALS and FTD.

## Materials and methods

### Protein Dataset Preparation

Starting structures for docking analysis and molecular dynamics simulations were extracted from the Protein Data Bank.^57^ We retrieved three TDP-43 crystal structures PDB ID: 7KWZ, 6N37, 7PY2).^46–48^ These include structures of ALS/FTLD patient-derived TDP-43 aggregates from the frontal and motor cortices, as well as two structures elucidated through crystallographic analysis. The selected PDB structures were prepared for computational approaches using the *Preparation wizard of Schrödinger*.^58^ The preparation of proteins includes adding hydrogens, ionizing metals, and fixing missing residues and atoms. To mimic the neutral pH conditions of the physiological environment, modifications were made such that the N-terminus was positively charged (NH3+) and the C-terminus was negatively charged (COO–). The prepared structures were used for docking and molecular dynamics analysis.

### Peptide Library Design

We generated a library of approximately 400 unique peptides, derived from the helical domain of TDP-43, with a length of 50 residues each, using the wildtype sequence [Uniprot (Q13148)]. Mutations were introduced at residue S332 and/or S333 in this region to generate 400 unique peptides. We utilized RaptorX (http://raptorx.uchicago.edu/StructurePropertyPred/predict/) to predict the secondary structure properties of the peptides.^52^ RaptorX Property is an advanced web server that can predict structural properties of a protein sequence without relying on any template information. This server excels in performance, particularly for proteins that lack closely related sequences in the Protein Data Bank (PDB) or have limited sequence profile information available. The peptides were then ranked based on their helical propensity, and the five top-ranked peptides with the highest helical propensity were selected for further analysis.

### Ab initio 3D Structure Prediction And Preparations

To validate sequence based secondary structural properties, the top five peptide hits with highest helical propensity were selected for 3D-structure prediction and further analysis. The top-five peptide structures were predicted using comparative modeling with the Rosetta comparative modeling (RosettaCM method).^53^ Wild-type TDP-43 amyloidogenic core region solution structure (PDB ID: 2N3X)^42^ was used as a template for the comparative modeling. This region spans 50 amino acids and encompasses residues 311-360 of the TDP-43 C-terminal LCD. The residues from 311-320 (*MNFGAFSINP*) and 344-360 (*QNQSGPSGNNQNQGNMQ*) are disorder coils, residues 321-330 (*AMMAAAQAAL*) and 335-343 (*GMMGMLASQ*) are known as conserved helical regions. The predicted peptides structures were used for further preparation, such as adding hydrogens, ionizing metals, and fixing missing atoms, with the help of the *preparation wizard from Schrödinger*.

### Docking Studies

Docking analysis was undertaken with the top five peptide candidates and amyloid-like filament structures (PDB ID: 7KWZ, 7PY2, 6N37) ***(Supplementary Fig.3B)***. This study was facilitated using the locally installed Hdock software package, following standard procedures. Based on the positions of peptides, docking scores, and favorable interactions with amyloid-like fibrils. We selected the top complex poses for molecular dynamics simulation.

### Molecular Dynamics Simulation

The top-ranked peptides and complexes were subjected to molecular dynamics simulations using GROMACS v2022. These simulations enable us to understand secondary structural properties, stability, intramolecular and intermolecular interactions, structural deviations, and fluctuations. Using the Optimized Potentials for Liquid Simulations (OPLS) force field, topologies for the leading complex were constructed.^55,59^ Using the Simple Point-Charge water model (SPCE), the system was placed in a cubic box configuration and solved.^60^ The energy was minimized until the steepest descent energy was attained, thereby aligning the atoms in order to reduce their maximum net stresses. This reduced the force applied to each atom, providing a favorable starting point for the simulations of molecular dynamics. Subsequently, elevated pressure and increasing temperature conditions were imposed for 100 ps, following which, the final molecular dynamics run generated a trajectory of 100 ns or 50 ns.

### Analysis Methods

The Root Mean Square Deviation (RMSD), Root Mean Square Fluctuation (RMSF), and secondary structural properties were calculated from trajectory data. To analyze the free energy landscape (FEL) and generate 3D figures, the following steps were employed. First, the structural coordinates, including RMSD, and Radius of Gyration (RG), helical probability and beta sheet probability, were extracted from trajectory data. Next, input files for Sham analysis were generated using the “sham.pl” Perl script. The Sham analysis was then performed using the “gmx sham” command, which produced the free energy landscape stored in the “free-energy-landscape.xpm” file. To further analyze, the “FEL.xpm” file was converted to a text format using the “xpm2txt.py” Python script. Finally, Mathematica 12.1, graphpad prism software were used to visualize the FEL, allowing for the generation of informative 3D figures. These methods provided a robust approach for investigating the free energy landscape and visualizing the results for comprehensive analysis. To predict the intramolecular interaction energy, we used Interaction energy matrix tool.^54^ Multiple structural conformations from various snapshots were retrieved from molecular dynamics trajectory data and used for predicting the intramolecular interaction energies.

## Author contribution

M.R.S and P.P.G designed the project, analyzed the results, and wrote the manuscript. M.R.S carried out the experiments.

## Acknowledgment

We thank past and present members of the Gopal lab for thoughtful discussion and comments on the manuscript.

