## Supplementary material for "Rational Design of TDP-43 Derived α-Helical Peptide Inhibitors: an *In-Silico* Strategy to Prevent TDP-43 Aggregation in Neurodegenerative Disorders": Salaikumaran_supporting_information_bioRxiv.docx

Supplementary Information

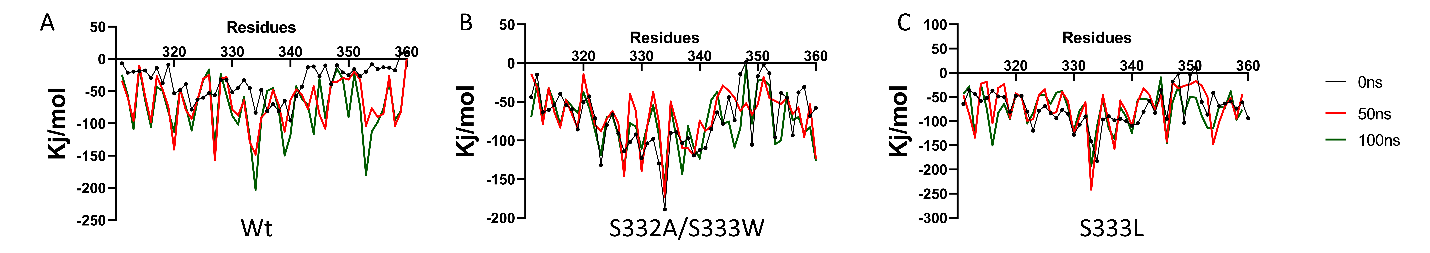

**Supplementary Figure 1: Calculated intra-molecular interaction energy (Kj/mol) at different timepoints(0, 50, 100 ns) for wild-type (Wt) (A) and helical propensity peptides**

**(B) S332/S333, (C) S333L.**

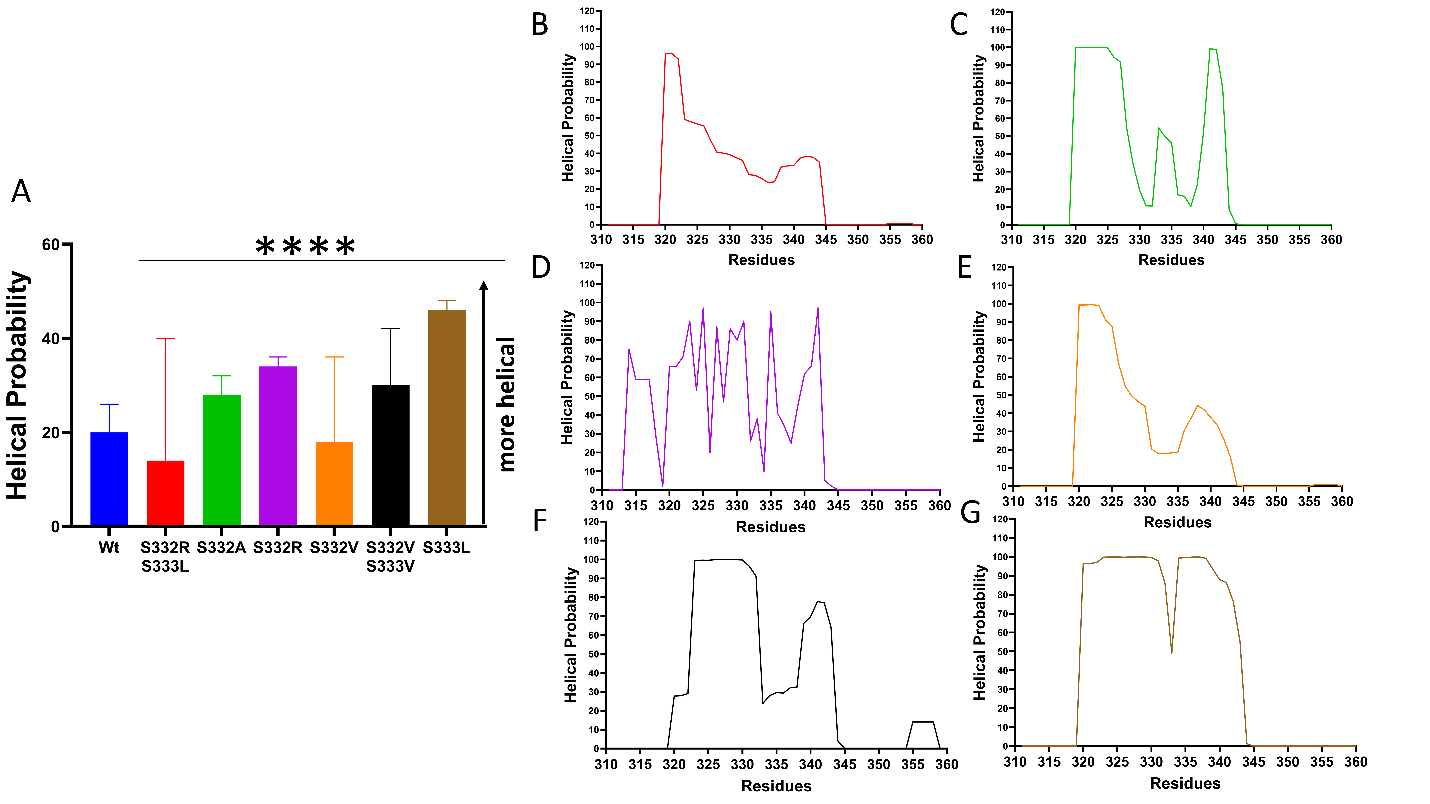

**Supplementary Figure 2: (A) Shows the overall helical probability for monomer residues 311-360 (P-value= <0.0001). (B-G): Shows the residues helical probability for S332RL(B), S332A(C), S332R(D) S332V(E), S332V/S333V(F), and S333V(G).**

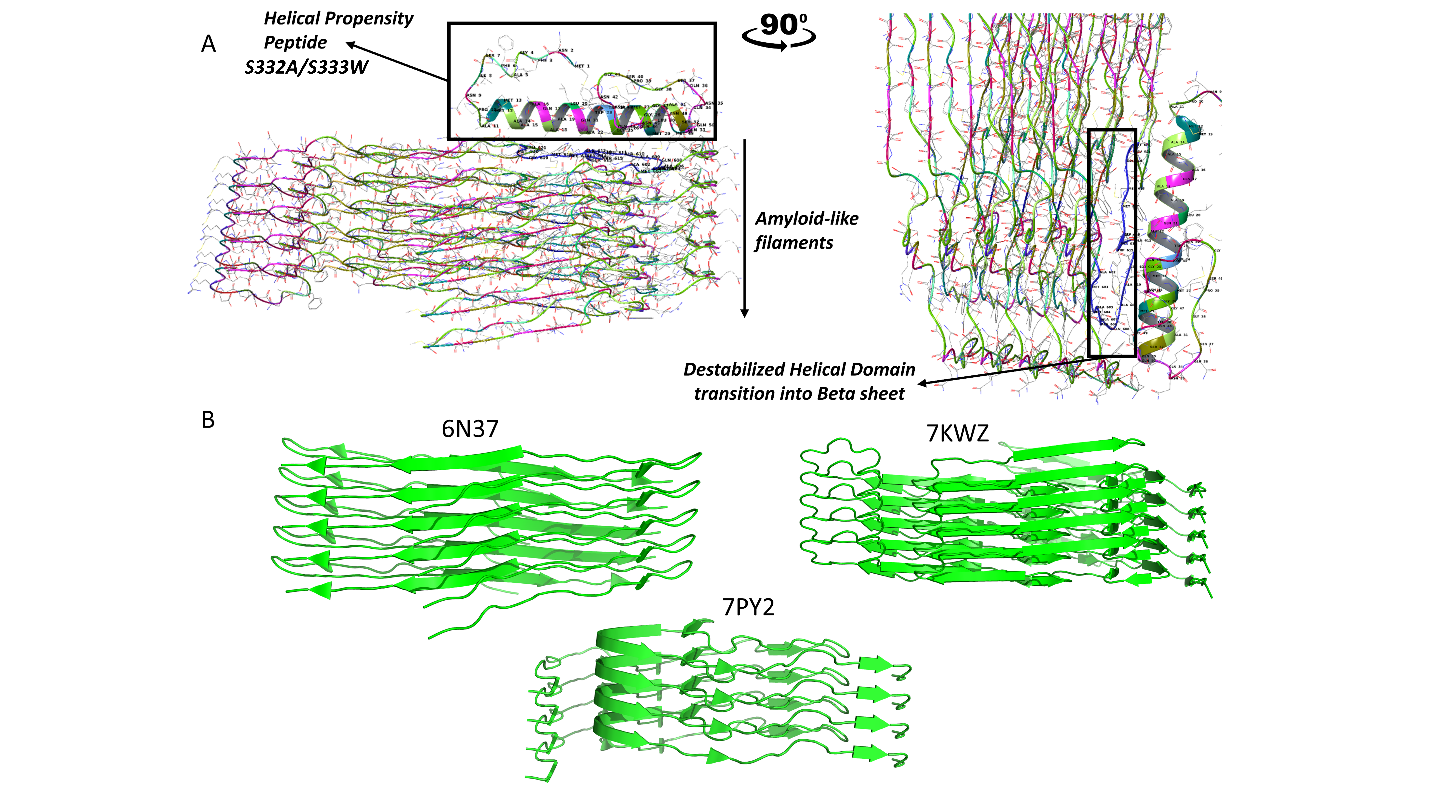

**Supplementary Figure 3: Molecular docking of the helical propensity variant S332A/S333W. A) Docking poses of S332A/S333W variant with the 7KWZ protein structure. B) Crystal structures employed in the molecular docking analysis.**

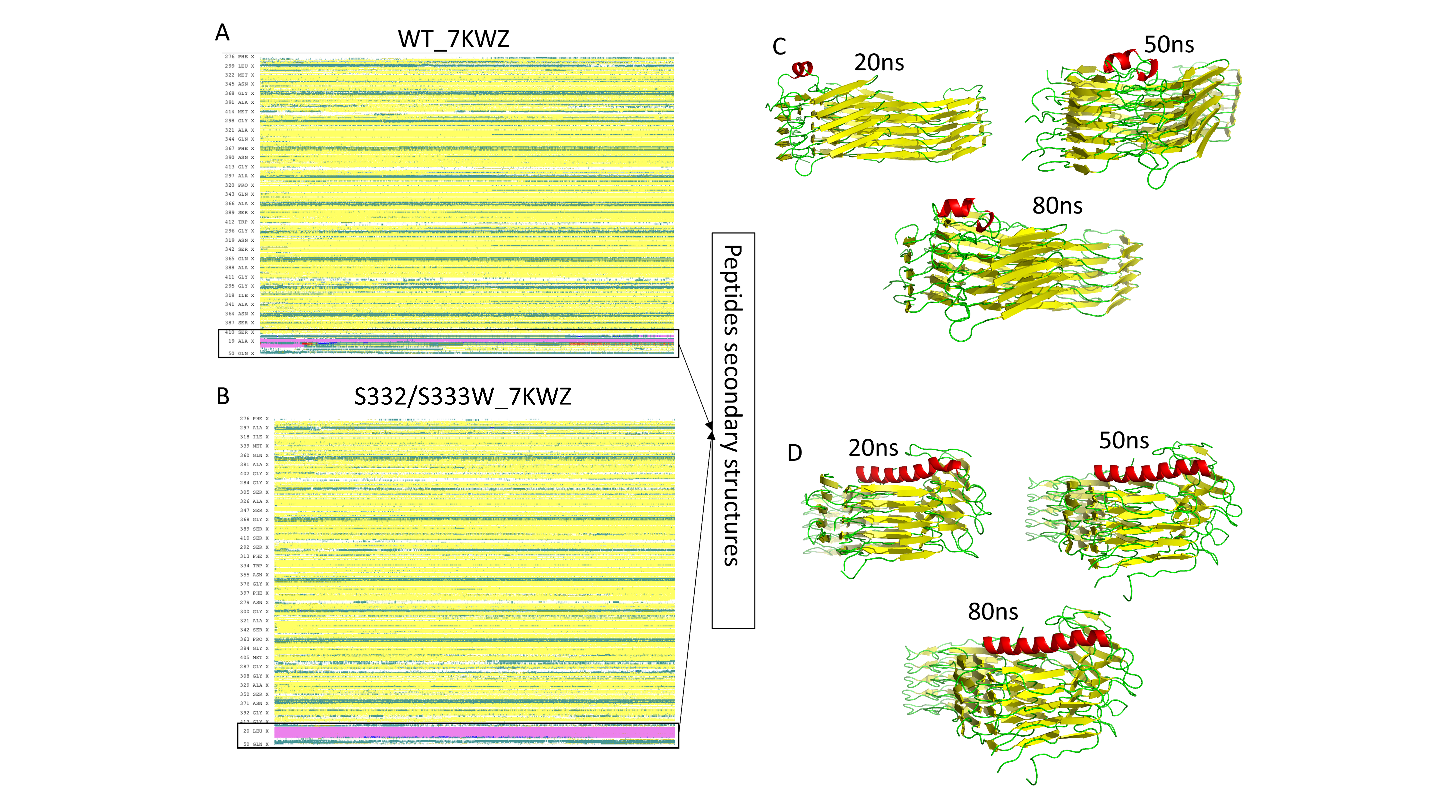

**Supplementary Figure 4: Structural and dynamic analysis of the wt and helical propensity variant complexes A) Predicted secondary structural properties of wt and helical propensity variants bound complexes over 100ns simulation period, displayed in distinct colors for each property(yellow-Beta sheets, green-loops, Pink-Helix). B) Structures of the variants bound with amyloid-like fibrils. C) Time-lapse snapshots from molecular dynamics simulations of wt and helical propensity variant-bound complexes.**

**Supplementary table.1: Predicted sequence based secondary structural properties for Serine 332 and Serine 333 mutant variants at Helical domain 320-340 TDP43**

| **Rank** | **Mutants** | **Sequence(320-340 Residues)** | **Helix Probability(320-340 Residues)**  **(H-helix, G-3Helix, T-turn, L-loop)** |
| --- | --- | --- | --- |
| 1 | >332_333AW | PAMMAAAQAALQAWWGMMGML | HHHHHHHHHHHHHGGGGGGGG |
| 2 | >332_333YC | PAMMAAAQAALQYCWGMMGML | HHHHHHHHHHHHHGGGGGGGG |
| 3 | >332_333RL | PAMMAAAQAALQRLWGMMGML | HHHHHHHHHHHHHGGGGGGGG |
| 4 | >332_333KL | PAMMAAAQAALQKLWGMMGML | HHHHHHHHHHHHGGGGGGGGG |
| 5 | >332_333YI | PAMMAAAQAALQYIWGMMGML | HHHHHHHHHHHHHHLGGGGGG |
| 6 | >332_333YL | PAMMAAAQAALQYLWGMMGML | HHHHHHHHHHHHHHLGGGGGG |
| 7 | >332_333IW | PAMMAAAQAALQIWWGMMGML | HHHHHHHHHHHHHHGGGGLGG |
| 8 | >332_333FR | PAMMAAAQAALQFRWGMMGML | HHHHHHHHHHHHHLGGGGGGG |
| 9 | >332_333YK | PAMMAAAQAALQYKWGMMGML | HHHHHHHHHHHHHLGGGGGGG |
| 10 | >332_333LR | PAMMAAAQAALQLRWGMMGML | HHHHHHHHHHHHHLGGGGGGG |
| 11 | >332_333VR | PAMMAAAQAALQVRWGMMGML | HHHHHHHHHHHHHLGGGGGGG |
| 12 | >332_333MG | PAMMAAAQAALQMGWGMMGML | HHHHHHHHHHHLGGGGGGGGG |
| 13 | >332_333WG | PAMMAAAQAALQWGWGMMGML | HHHHHHHHHHHLGGGGGGGGG |
| 14 | >332_333VL | PAMMAAAQAALQVLWGMMGML | HHHHHHHHHHHHHHGLGGLGG |
| 15 | >332_333VW | PAMMAAAQAALQVWWGMMGML | HHHHHHHHHHHHHHGGGGLLG |
| 16 | >332_333TA | PAMMAAAQAALQTAWGMMGML | HHHHHHHHHHHHHLGGGGLGG |
| 17 | >332_333TC | PAMMAAAQAALQTCWGMMGML | HHHHHHHHHHHHHLLGGGGGG |
| 18 | >332_333YE | PAMMAAAQAALQYEWGMMGML | HHHHHHHHHHHHHLLGGGGGG |
| 19 | >332_333FC | PAMMAAAQAALQFCWGMMGML | HHHHHHHHHHHHHLLGGGGGG |
| 20 | >332_333FM | PAMMAAAQAALQFMWGMMGML | HHHHHHHHHHHHHLLGGGGGG |
| 21 | >332_333YV | PAMMAAAQAALQYVWGMMGML | HHHHHHHHHHHHHLLGGGGGG |
| 22 | >332_333VC | PAMMAAAQAALQVCWGMMGML | HHHHHHHHHHHHHLLGGGGGG |
| 23 | >332_333CW | PAMMAAAQAALQCWWGMMGML | HHHHHHHHHHHHHLGGGGLGG |
| 24 | >332_333YW | PAMMAAAQAALQYWWGMMGML | HHHHHHHHHHHHHLLGGGGGG |
| 25 | >332_333TW | PAMMAAAQAALQTWWGMMGML | HHHHHHHHHHHHHLGGGGLGG |
| 26 | >332_333YR | PAMMAAAQAALQYRWGMMGML | HHHHHHHHHHHHHLLGGGGGG |
| 27 | >332_333YT | PAMMAAAQAALQYTWGMMGML | HHHHHHHHHHHHHLLGGGGGG |
| 28 | >332_333CA | PAMMAAAQAALQCAWGMMGML | HHHHHHHHHHHHLLGGGGGGG |
| 29 | >332_333FD | PAMMAAAQAALQFDWGMMGML | HHHHHHHHHHHHLLGGGGGGG |
| 30 | >332_333GF | PAMMAAAQAALQGFWGMMGML | HHHHHHHHHHHHLLGGGGGGG |
| 31 | >332_333GC | PAMMAAAQAALQGCWGMMGML | HHHHHHHHHHHHLLGGGGGGG |
| 32 | >332_333CG | PAMMAAAQAALQCGWGMMGML | HHHHHHHHHHHHLLGGGGGGG |
| 33 | >332_333VP | PAMMAAAQAALQVPWGMMGML | HHHHHHHHHHHHLLGGGGGGG |
| 34 | >332_333CP | PAMMAAAQAALQCPWGMMGML | HHHHHHHHHHHHLLGGGGGGG |
| 35 | >332_333YP | PAMMAAAQAALQYPWGMMGML | HHHHHHHHHHHHLLGGGGGGG |
| 36 | >332_333GM | PAMMAAAQAALQGMWGMMGML | HHHHHHHHHHHLLGGGGGGGG |
| 37 | >332_333IG | PAMMAAAQAALQIGWGMMGML | HHHHHHHHHHHLLGGGGGGGG |
| 38 | >332_333TG | PAMMAAAQAALQTGWGMMGML | HHHHHHHHHHHLLGGGGGGGG |
| 39 | >332_333WF | PAMMAAAQAALQWFWGMMGML | HHHHHHHHHHHHHHGGGGLLL |
| 40 | >332_333LW | PAMMAAAQAALQLWWGMMGML | HHHHHHHHHHHHHHGGGGLTL |
| 41 | >332_333WQ | PAMMAAAQAALQWQWGMMGML | HHHHHHHHHHHHHHGGGGLLL |
| 42 | >332_333WY | PAMMAAAQAALQWYWGMMGML | HHHHHHHHHHHHHHGGGGLLL |
| 43 | >332_333RW | PAMMAAAQAALQRWWGMMGML | HHHHHHHHHHHHHHGGGGLTL |
| 44 | >332_333SF | PAMMAAAQAALQSFWGMMGML | HHHHHHHHHHHHHTTLGGGGG |
| 45 | >332_333AC | PAMMAAAQAALQACWGMMGML | HHHHHHHHHHHHHTLLGGGGG |
| 46 | >332_333AY | PAMMAAAQAALQAYWGMMGML | HHHHHHHHHHHHHTLLGGGGG |
| 47 | >332_333IC | PAMMAAAQAALQICWGMMGML | HHHHHHHHHHHHHLLGGGLGG |
| 48 | >332_333YM | PAMMAAAQAALQYMWGMMGML | HHHHHHHHHHHHHLLLGGGGG |
| 49 | >332_333QR | PAMMAAAQAALQQRWGMMGML | HHHHHHHHHHHHHLLLGGGGG |
| 50 | >332_333YS | PAMMAAAQAALQYSWGMMGML | HHHHHHHHHHHHLLLGGGGGG |
| 51 | >332_333CR | PAMMAAAQAALQCRWGMMGML | HHHHHHHHHHHHLLLGGGGGG |
| 52 | >332_333CE | PAMMAAAQAALQCEWGMMGML | HHHHHHHHHHHHLLLGGGGGG |
| 53 | >332_333GL | PAMMAAAQAALQGLWGMMGML | HHHHHHHHHHHHLLGGGGLGG |
| 54 | >332_333YG | PAMMAAAQAALQYGWGMMGML | HHHHHHHHHHHHLLLGGGGGG |
| 55 | >332_333KC | PAMMAAAQAALQKCWGMMGML | HHHHHHHHHHHHLLLGGGGGG |
| 56 | >332_333YQ | PAMMAAAQAALQYQWGMMGML | HHHHHHHHHHHHTLLGGGGGG |
| 57 | >332_333WW | PAMMAAAQAALQWWWGMMGML | GGHHHHHHHHHHHHGGGGLLL |
| 58 | >332_333YN | PAMMAAAQAALQYNWGMMGML | HHHHHHHHHHHHTLLGGGGGG |
| 59 | >332_333YD | PAMMAAAQAALQYDWGMMGML | HHHHHHHHHHHHLLLGGGGGG |
| 60 | >332_333AP | PAMMAAAQAALQAPWGMMGML | HHHHHHHHHHHLLLGGGGGGG |
| 61 | >332_333EP | PAMMAAAQAALQEPWGMMGML | HHHHHHHHHHHLLLGGGGGGG |
| 62 | >332_333VG | PAMMAAAQAALQVGWGMMGML | HHHHHHHHHHHLLLGGGGGGG |
| 63 | >332_333KP | PAMMAAAQAALQKPWGMMGML | HHHHHHHHHHHLLLGGGGGGG |
| 64 | >332_333PA | PAMMAAAQAALQPAWGMMGML | HHHHHHHHHHLLLGGGGGGGG |
| 65 | >332_333PF | PAMMAAAQAALQPFWGMMGML | HHHHHHHHHHLLLGGGGGGGG |
| 66 | >332_333PK | PAMMAAAQAALQPKWGMMGML | HHHHHHHHHHLLLGGGGGGGG |
| 67 | >332_333ND | PAMMAAAQAALQNDWGMMGML | HHHHHHHHHHLLLGGGGGGGG |
| 68 | >332_333RP | PAMMAAAQAALQRPWGMMGML | HHHHHHHHHHLLLGGGGGGGG |
| 69 | >332_333PY | PAMMAAAQAALQPYWGMMGML | HHHHHHHHHHLLLGGGGGGGG |
| 70 | >332_333PD | PAMMAAAQAALQPDWGMMGML | HHHHHHHHHHLLLGGGGGGGG |
| 71 | >332_333PW | PAMMAAAQAALQPWWGMMGML | HHHHHHHHHHLLLGGGGGGGG |
| 72 | >332_333AI | PAMMAAAQAALQAIWGMMGML | HHHHHHHHHHHHHHGLLLLGG |
| 73 | >332_333RV | PAMMAAAQAALQRVWGMMGML | HHHHHHHHHHHHHHGGGLLLL |
| 74 | >332_333EW | PAMMAAAQAALQEWWGMMGML | HHHHHHHHHHHHHHGGGLLLL |
| 75 | >332_333CM | PAMMAAAQAALQCMWGMMGML | HHHHHHHHHHHHHLLGGGLLG |
| 76 | >332_333GV | PAMMAAAQAALQGVWGMMGML | HHHHHHHHHHHHLLLLGGGGG |
| 77 | >332_333GY | PAMMAAAQAALQGYWGMMGML | HHHHHHHHHHHHTLLGGGLGG |
| 78 | >332_333IK | PAMMAAAQAALQIKWGMMGML | HHHHHHHHHHHHLLGGGGLLG |
| 79 | >332_333CQ | PAMMAAAQAALQCQWGMMGML | HHHHHHHHHHHHLLLLGGGGG |
| 80 | >332_333FG | PAMMAAAQAALQFGWGMMGML | HHHHHHHHHHHLLLLGGGGGG |
| 81 | >332_333WP | PAMMAAAQAALQWPWGMMGML | HHHHHHHHHHHLLLGGGGLGG |
| 82 | >332_333SP | PAMMAAAQAALQSPWGMMGML | HHHHHHHHHHLLLTGGGGGGG |
| 83 | >332_333PS | PAMMAAAQAALQPSWGMMGML | HHHHHHHHHHLLLLGGGGGGG |
| 84 | >332_333NE | PAMMAAAQAALQNEWGMMGML | HHHHHHHHHHLLLLGGGGGGG |
| 85 | >332_333PP | PAMMAAAQAALQPPWGMMGML | HHHHHHHHHHLLLLGGGGGGG |
| 86 | >332_333TP | PAMMAAAQAALQTPWGMMGML | HHHHHHHHHHLLLLGGGGGGG |
| 87 | >332_333PI | PAMMAAAQAALQPIWGMMGML | HHHHHHHHHHLLLLGGGGGGG |
| 88 | >332_333PC | PAMMAAAQAALQPCWGMMGML | HHHHHHHHHHLLLLGGGGGGG |
| 89 | >332_333PR | PAMMAAAQAALQPRWGMMGML | HHHHHHHHHHLLLLGGGGGGG |
| 90 | >332_333PN | PAMMAAAQAALQPNWGMMGML | HHHHHHHHHHLLLLGGGGGGG |
| 91 | >332_333PT | PAMMAAAQAALQPTWGMMGML | HHHHHHHHHHLLLLGGGGGGG |
| 92 | >332_333PQ | PAMMAAAQAALQPQWGMMGML | HHHHHHHHHHLLLLGGGGGGG |
| 93 | >332_333PE | PAMMAAAQAALQPEWGMMGML | HHHHHHHHHLLLLGGGGGGGG |
| 94 | >332_333PG | PAMMAAAQAALQPGWGMMGML | HHHHHHHHHLLLLGGGGGGGG |
| 95 | >332_333EV | PAMMAAAQAALQEVWGMMGML | HHHHHHHHHHHHHHHLGLLLL |
| 96 | >332_333EL | PAMMAAAQAALQELWGMMGML | HHHHHHHHHHHHHHHLGTLLL |
| 97 | >332_333AM | PAMMAAAQAALQAMWGMMGML | HHHHHHHHHHHHHHLLLLLGG |
| 98 | >332_333VA | PAMMAAAQAALQVAWGMMGML | HHHHHHHHHHHHHHLLLLLGG |
| 99 | >332_333LQ | PAMMAAAQAALQLQWGMMGML | HHHHHHHHHHHHHHLLLLLGG |
| 100 | >332_333WC | PAMMAAAQAALQWCWGMMGML | HHHHHHHHHHHHHHLLGGLLL |
| 101 | >332_333DW | PAMMAAAQAALQDWWGMMGML | HHHHHHHHHHHHHHLLGGLLL |
| 102 | >332_333TY | PAMMAAAQAALQTYWGMMGML | HHHHHHHHHHHHHHLLGGLLL |
| 103 | >332_333RA | PAMMAAAQAALQRAWGMMGML | HHHHHHHHHHHHHLGLLLLGG |
| 104 | >332_333CC | PAMMAAAQAALQCCWGMMGML | HHHHHHHHHHHHHLLLGGGLL |
| 105 | >332_333IR | PAMMAAAQAALQIRWGMMGML | HHHHHHHHHHHHHLLGGGLLL |
| 106 | >332_333LC | PAMMAAAQAALQLCWGMMGML | HHHHHHHHHHHHHLLLGGLGL |
| 107 | >332_333MC | PAMMAAAQAALQMCWGMMGML | HHHHHHHHHHHHHLLLLLGGG |
| 108 | >332_333RC | PAMMAAAQAALQRCWGMMGML | HHHHHHHHHHHHLLLGGGLLG |
| 109 | >332_333CN | PAMMAAAQAALQCNWGMMGML | HHHHHHHHHHHHLLLLGGLGG |
| 110 | >332_333VE | PAMMAAAQAALQVEWGMMGML | HHHHHHHHHHHHLLLLGGLGG |
| 111 | >332_333CF | PAMMAAAQAALQCFWGMMGML | HHHHHHHHHHHHLLLLGGLGG |
| 112 | >332_333CK | PAMMAAAQAALQCKWGMMGML | HHHHHHHHHHHHTLLLGGLGG |
| 113 | >332_333HP | PAMMAAAQAALQHPWGMMGML | HHHHHHHHHHLLLTTGGGGGG |
| 114 | >332_333DP | PAMMAAAQAALQDPWGMMGML | HHHHHHHHHHLLLTTGGGGGG |
| 115 | >332_333PV | PAMMAAAQAALQPVWGMMGML | HHHHHHHHHLLLLLGGGGGGG |
| 116 | >332_333LA | PAMMAAAQAALQLAWGMMGML | HHHHHHHHHHHHHHHLLLLLL |
| 117 | >332_333WA | PAMMAAAQAALQWAWGMMGML | HHHHHHHHHHHHHHHLLLLLL |
| 118 | >332_333EA | PAMMAAAQAALQEAWGMMGML | HHHHHHHHHHHHHHHLLLLLL |
| 119 | >332_333YA | PAMMAAAQAALQYAWGMMGML | HHHHHHHHHHHHHHHLLLLLL |
| 120 | >332_333II | PAMMAAAQAALQIIWGMMGML | HHHHHHHHHHHHHHHLLLLLL |
| 121 | >332_333CI | PAMMAAAQAALQCIWGMMGML | HHHHHHHHHHHHHHHLLLLLL |
| 122 | >332_333LI | PAMMAAAQAALQLIWGMMGML | HHHHHHHHHHHHHHHLLLLLL |
| 123 | >332_333ML | PAMMAAAQAALQMLWGMMGML | HHHHHHHHHHHHHHHLLLLLL |
| 124 | >332_333WL | PAMMAAAQAALQWLWGMMGML | HHHHHHHHHHHHHHHLLLLLL |
| 125 | >332_333IL | PAMMAAAQAALQILWGMMGML | HHHHHHHHHHHHHHHLLLLLL |
| 126 | >332_333CL | PAMMAAAQAALQCLWGMMGML | HHHHHHHHHHHHHHHLLLLLL |
| 127 | >332_333QI | PAMMAAAQAALQQIWGMMGML | HHHHHHHHHHHHHHHLLLLLL |
| 128 | >332_333VV | PAMMAAAQAALQVVWGMMGML | HHHHHHHHHHHHHHHLLLLLL |
| 129 | >332_333CV | PAMMAAAQAALQCVWGMMGML | HHHHHHHHHHHHHHHLLLLLL |
| 130 | >332_333VI | PAMMAAAQAALQVIWGMMGML | HHHHHHHHHHHHHHHLLLLLL |
| 131 | >332_333WE | PAMMAAAQAALQWEWGMMGML | HHHHHHHHHHHHHHHLLLLLL |
| 132 | >332_333WV | PAMMAAAQAALQWVWGMMGML | HHHHHHHHHHHHHHHLLLLLL |
| 133 | >332_333WI | PAMMAAAQAALQWIWGMMGML | HHHHHHHHHHHHHHHLLLLLL |
| 134 | >332_333WH | PAMMAAAQAALQWHWGMMGML | HHHHHHHHHHHHHHHLLLLLL |
| 135 | >332_333AT | PAMMAAAQAALQATWGMMGML | HHHHHHHHHHHHHHLLLLLLG |
| 136 | >332_333QW | PAMMAAAQAALQQWWGMMGML | HHHHHHHHHHHHHHLLGLLLL |
| 137 | >332_333NC | PAMMAAAQAALQNCWGMMGML | HHHHHHHHHHHHHLLLGGLLL |
| 138 | >332_333FW | PAMMAAAQAALQFWWGMMGML | HHHHHHHHHHHHHLGLGLLLL |
| 139 | >332_333FN | PAMMAAAQAALQFNWGMMGML | HHHHHHHHHHHHHLLGGLLLL |
| 140 | >332_333NL | PAMMAAAQAALQNLWGMMGML | HHHHHHHHHHHHHLLGGLLLL |
| 141 | >332_333RR | PAMMAAAQAALQRRWGMMGML | HHHHHHHHHHHHHLLLGGLLL |
| 142 | >332_333RN | PAMMAAAQAALQRNWGMMGML | HHHHHHHHHHHHHLLLGGLLL |
| 143 | >332_333RD | PAMMAAAQAALQRDWGMMGML | HHHHHHHHHHHHHLLLGGLLL |
| 144 | >332_333SW | PAMMAAAQAALQSWWGMMGML | HHHHHHHHHHHHLLGGGLLLL |
| 145 | >332_333CT | PAMMAAAQAALQCTWGMMGML | HHHHHHHHHHHHLLLLGLLGG |
| 146 | >332_333TI | PAMMAAAQAALQTIWGMMGML | HHHHHHHHHHHHLLLGGLLLG |
| 147 | >332_333LN | PAMMAAAQAALQLNWGMMGML | HHHHHHHHHHHHLLLLLLGGG |
| 148 | >332_333AA | PAMMAAAQAALQAAWGMMGML | HHHHHHHHHHHHHHLLLLLLL |
| 149 | >332_333AL | PAMMAAAQAALQALWGMMGML | HHHHHHHHHHHHHHLLLLLLL |
| 150 | >332_333AQ | PAMMAAAQAALQAQWGMMGML | HHHHHHHHHHHHHHLLLLLLL |
| 151 | >332_333AE | PAMMAAAQAALQAEWGMMGML | HHHHHHHHHHHHHHLLLLLLL |
| 152 | >332_333AV | PAMMAAAQAALQAVWGMMGML | HHHHHHHHHHHHHHLLLLLLL |
| 153 | >332_333KA | PAMMAAAQAALQKAWGMMGML | HHHHHHHHHHHHHHLLLLLLL |
| 154 | >332_333QA | PAMMAAAQAALQQAWGMMGML | HHHHHHHHHHHHHHLLLLLLL |
| 155 | >332_333IA | PAMMAAAQAALQIAWGMMGML | HHHHHHHHHHHHHHLLLLLLL |
| 156 | >332_333HA | PAMMAAAQAALQHAWGMMGML | HHHHHHHHHHHHHHLLLLLLL |
| 157 | >332_333DA | PAMMAAAQAALQDAWGMMGML | HHHHHHHHHHHHHHLLLLLLL |
| 158 | >332_333EH | PAMMAAAQAALQEHWGMMGML | HHHHHHHHHHHHHHLLLLLLL |
| 159 | >332_333EF | PAMMAAAQAALQEFWGMMGML | HHHHHHHHHHHHHHLLLLLLL |
| 160 | >332_333RI | PAMMAAAQAALQRIWGMMGML | HHHHHHHHHHHHHHLLLLLLL |
| 161 | >332_333IY | PAMMAAAQAALQIYWGMMGML | HHHHHHHHHHHHHHLLLLLLL |
| 162 | >332_333KI | PAMMAAAQAALQKIWGMMGML | HHHHHHHHHHHHHHLLLLLLL |
| 163 | >332_333LL | PAMMAAAQAALQLLWGMMGML | HHHHHHHHHHHHHHLLLLLLL |
| 164 | >332_333LV | PAMMAAAQAALQLVWGMMGML | HHHHHHHHHHHHHHLLLLLLL |
| 165 | >332_333LY | PAMMAAAQAALQLYWGMMGML | HHHHHHHHHHHHHHLLLLLLL |
| 166 | >332_333FL | PAMMAAAQAALQFLWGMMGML | HHHHHHHHHHHHHHLLLLLLL |
| 167 | >332_333QL | PAMMAAAQAALQQLWGMMGML | HHHHHHHHHHHHHHLLLLLLL |
| 168 | >332_333MI | PAMMAAAQAALQMIWGMMGML | HHHHHHHHHHHHHHLLLLLLL |
| 169 | >332_333WM | PAMMAAAQAALQWMWGMMGML | HHHHHHHHHHHHHHLLLLLLL |
| 170 | >332_333IM | PAMMAAAQAALQIMWGMMGML | HHHHHHHHHHHHHHLLLLLLL |
| 171 | >332_333IV | PAMMAAAQAALQIVWGMMGML | HHHHHHHHHHHHHHLLLLLLL |
| 172 | >332_333WK | PAMMAAAQAALQWKWGMMGML | HHHHHHHHHHHHHHTLLLLLL |
| 173 | >332_333WR | PAMMAAAQAALQWRWGMMGML | HHHHHHHHHHHHHHTLLLLLL |
| 174 | >332_333WT | PAMMAAAQAALQWTWGMMGML | HHHHHHHHHHHHHHLLLLLLL |
| 175 | >332_333CH | PAMMAAAQAALQCHWGMMGML | HHHHHHHHHHHHHLLLGLLLL |
| 176 | >332_333FF | PAMMAAAQAALQFFWGMMGML | HHHHHHHHHHHHHLLLLLLLG |
| 177 | >332_333IF | PAMMAAAQAALQIFWGMMGML | HHHHHHHHHHHHHLLLGLLLL |
| 178 | >332_333YF | PAMMAAAQAALQYFWGMMGML | HHHHHHHHHHHHHLLLLLLLG |
| 179 | >332_333HF | PAMMAAAQAALQHFWGMMGML | HHHHHHHHHHHHHLLLGLLLL |
| 180 | >332_333IH | PAMMAAAQAALQIHWGMMGML | HHHHHHHHHHHHHTLLGLLLL |
| 181 | >332_333LD | PAMMAAAQAALQLDWGMMGML | HHHHHHHHHHHHHLLLLGLLL |
| 182 | >332_333TL | PAMMAAAQAALQTLWGMMGML | HHHHHHHHHHHHHLLLLLLLG |
| 183 | >332_333QC | PAMMAAAQAALQQCWGMMGML | HHHHHHHHHHHHHLLLGLLLL |
| 184 | >332_333VH | PAMMAAAQAALQVHWGMMGML | HHHHHHHHHHHHHLLLGLLLL |
| 185 | >332_333VD | PAMMAAAQAALQVDWGMMGML | HHHHHHHHHHHHHLLLLLLLG |
| 186 | >332_333HW | PAMMAAAQAALQHWWGMMGML | HHHHHHHHHHHHHLLLGLLLL |
| 187 | >332_333DC | PAMMAAAQAALQDCWGMMGML | HHHHHHHHHHHHLLLLGGLLL |
| 188 | >332_333CY | PAMMAAAQAALQCYWGMMGML | HHHHHHHHHHHHLLLLGLLLG |
| 189 | >332_333CD | PAMMAAAQAALQCDWGMMGML | HHHHHHHHHHHHLLLLGGLLL |
| 190 | >332_333IE | PAMMAAAQAALQIEWGMMGML | HHHHHHHHHHHHLLLLLLLGG |
| 191 | >332_333GI | PAMMAAAQAALQGIWGMMGML | HHHHHHHHHHHHTLLGGLLLL |
| 192 | >332_333LK | PAMMAAAQAALQLKWGMMGML | HHHHHHHHHHHHLLLLLLLGG |
| 193 | >332_333WD | PAMMAAAQAALQWDWGMMGML | HHHHHHHHHHHHLLLLGGLLL |
| 194 | >332_333FP | PAMMAAAQAALQFPWGMMGML | HHHHHHHHHHHLLLTLGGLLG |
| 195 | >332_333DG | PAMMAAAQAALQDGWGMMGML | HHHHHHHHHHHLLLLGGGLLL |
| 196 | >332_333QN | PAMMAAAQAALQQNWGMMGML | HHHHHHHHHHHLLLLGGGLLL |
| 197 | >332_333IP | PAMMAAAQAALQIPWGMMGML | HHHHHHHHHHLLLLTGGGLLG |
| 198 | >332_333SA | PAMMAAAQAALQSAWGMMGML | HHHHHHHHHHHHHLLLLLLLL |
| 199 | >332_333SM | PAMMAAAQAALQSMWGMMGML | HHHHHHHHHHHHHLLLLLLLL |
| 200 | >332_333SC | PAMMAAAQAALQSCWGMMGML | HHHHHHHHHHHHHLLLLLLLL |
| 201 | >332_333SY | PAMMAAAQAALQSYWGMMGML | HHHHHHHHHHHHHLLLLLLLL |
| 202 | >332_333SH | PAMMAAAQAALQSHWGMMGML | HHHHHHHHHHHHHLLLLLLLL |
| 203 | >332_333SR | PAMMAAAQAALQSRWGMMGML | HHHHHHHHHHHHHLLLLLLLL |
| 204 | >332_333ST | PAMMAAAQAALQSTWGMMGML | HHHHHHHHHHHHHLLLLLLLL |
| 205 | >332_333WS | PAMMAAAQAALQWSWGMMGML | HHHHHHHHHHHHHLLLLLLLL |
| 206 | >332_333KS | PAMMAAAQAALQKSWGMMGML | HHHHHHHHHHHHHLLLLLLLL |
| 207 | >332_333ES | PAMMAAAQAALQESWGMMGML | HHHHHHHHHHHHHLLLLLLLL |
| 208 | >332_333RS | PAMMAAAQAALQRSWGMMGML | HHHHHHHHHHHHHLLLLLLLL |
| 209 | >332_333TS | PAMMAAAQAALQTSWGMMGML | HHHHHHHHHHHHHLLLLLLLL |
| 210 | >332_333AF | PAMMAAAQAALQAFWGMMGML | HHHHHHHHHHHHHTLLLLLLL |
| 211 | >332_333AK | PAMMAAAQAALQAKWGMMGML | HHHHHHHHHHHHHTLLLLLLL |
| 212 | >332_333AH | PAMMAAAQAALQAHWGMMGML | HHHHHHHHHHHHHTLLLLLLL |
| 213 | >332_333AR | PAMMAAAQAALQARWGMMGML | HHHHHHHHHHHHHTLLLLLLL |
| 214 | >332_333AN | PAMMAAAQAALQANWGMMGML | HHHHHHHHHHHHHLLLLLLLL |
| 215 | >332_333AD | PAMMAAAQAALQADWGMMGML | HHHHHHHHHHHHHLLLLLLLL |
| 216 | >332_333MA | PAMMAAAQAALQMAWGMMGML | HHHHHHHHHHHHHLLLLLLLL |
| 217 | >332_333FA | PAMMAAAQAALQFAWGMMGML | HHHHHHHHHHHHHLLLLLLLL |
| 218 | >332_333NA | PAMMAAAQAALQNAWGMMGML | HHHHHHHHHHHHHLLLLLLLL |
| 219 | >332_333HC | PAMMAAAQAALQHCWGMMGML | HHHHHHHHHHHHHLLLLLLLL |
| 220 | >332_333TT | PAMMAAAQAALQTTWGMMGML | HHHHHHHHHHHHHLLLLLLLL |
| 221 | >332_333DT | PAMMAAAQAALQDTWGMMGML | HHHHHHHHHHHHHLLLLLLLL |
| 222 | >332_333EE | PAMMAAAQAALQEEWGMMGML | HHHHHHHHHHHHHLLLLLLLL |
| 223 | >332_333HE | PAMMAAAQAALQHEWGMMGML | HHHHHHHHHHHHHLLLLLLLL |
| 224 | >332_333RE | PAMMAAAQAALQREWGMMGML | HHHHHHHHHHHHHLLLLLLLL |
| 225 | >332_333EI | PAMMAAAQAALQEIWGMMGML | HHHHHHHHHHHHHLLLLLLLL |
| 226 | >332_333EC | PAMMAAAQAALQECWGMMGML | HHHHHHHHHHHHHLLLLLLLL |
| 227 | >332_333ER | PAMMAAAQAALQERWGMMGML | HHHHHHHHHHHHHLLLLLLLL |
| 228 | >332_333EN | PAMMAAAQAALQENWGMMGML | HHHHHHHHHHHHHLLLLLLLL |
| 229 | >332_333ET | PAMMAAAQAALQETWGMMGML | HHHHHHHHHHHHHTLLLLLLL |
| 230 | >332_333FQ | PAMMAAAQAALQFQWGMMGML | HHHHHHHHHHHHHLLLLLLLL |
| 231 | >332_333FV | PAMMAAAQAALQFVWGMMGML | HHHHHHHHHHHHHLLLLLLLL |
| 232 | >332_333FY | PAMMAAAQAALQFYWGMMGML | HHHHHHHHHHHHHLLLLLLLL |
| 233 | >332_333FH | PAMMAAAQAALQFHWGMMGML | HHHHHHHHHHHHHLLLLLLLL |
| 234 | >332_333KF | PAMMAAAQAALQKFWGMMGML | HHHHHHHHHHHHHLLLLLLLL |
| 235 | >332_333QF | PAMMAAAQAALQQFWGMMGML | HHHHHHHHHHHHHLLLLLLLL |
| 236 | >332_333VF | PAMMAAAQAALQVFWGMMGML | HHHHHHHHHHHHHLLLLLLLL |
| 237 | >332_333RF | PAMMAAAQAALQRFWGMMGML | HHHHHHHHHHHHHLLLLLLLL |
| 238 | >332_333TF | PAMMAAAQAALQTFWGMMGML | HHHHHHHHHHHHHLLLLLLLL |
| 239 | >332_333HH | PAMMAAAQAALQHHWGMMGML | HHHHHHHHHHHHHLLLLLLLL |
| 240 | >332_333RH | PAMMAAAQAALQRHWGMMGML | HHHHHHHHHHHHHTLLLLLLL |
| 241 | >332_333NH | PAMMAAAQAALQNHWGMMGML | HHHHHHHHHHHHHTLLLLLLL |
| 242 | >332_333DH | PAMMAAAQAALQDHWGMMGML | HHHHHHHHHHHHHLLLLLLLL |
| 243 | >332_333TH | PAMMAAAQAALQTHWGMMGML | HHHHHHHHHHHHHLLLLLLLL |
| 244 | >332_333HR | PAMMAAAQAALQHRWGMMGML | HHHHHHHHHHHHHLLLLLLLL |
| 245 | >332_333HN | PAMMAAAQAALQHNWGMMGML | HHHHHHHHHHHHHLLLLLLLL |
| 246 | >332_333HT | PAMMAAAQAALQHTWGMMGML | HHHHHHHHHHHHHLLLLLLLL |
| 247 | >332_333NI | PAMMAAAQAALQNIWGMMGML | HHHHHHHHHHHHHLLLLLLLL |
| 248 | >332_333ID | PAMMAAAQAALQIDWGMMGML | HHHHHHHHHHHHHLLLLLLLL |
| 249 | >332_333KK | PAMMAAAQAALQKKWGMMGML | HHHHHHHHHHHHHLLLLLLLL |
| 250 | >332_333KQ | PAMMAAAQAALQKQWGMMGML | HHHHHHHHHHHHHLLLLLLLL |
| 251 | >332_333KE | PAMMAAAQAALQKEWGMMGML | HHHHHHHHHHHHHLLLLLLLL |
| 252 | >332_333KV | PAMMAAAQAALQKVWGMMGML | HHHHHHHHHHHHHLLLLLLLL |
| 253 | >332_333KY | PAMMAAAQAALQKYWGMMGML | HHHHHHHHHHHHHTLLLLLLL |
| 254 | >332_333KH | PAMMAAAQAALQKHWGMMGML | HHHHHHHHHHHHHLLLLLLLL |
| 255 | >332_333KR | PAMMAAAQAALQKRWGMMGML | HHHHHHHHHHHHHLLLLLLLL |
| 256 | >332_333KN | PAMMAAAQAALQKNWGMMGML | HHHHHHHHHHHHHLLLLLLLL |
| 257 | >332_333KT | PAMMAAAQAALQKTWGMMGML | HHHHHHHHHHHHHLLLLLLLL |
| 258 | >332_333EK | PAMMAAAQAALQEKWGMMGML | HHHHHHHHHHHHHLLLLLLLL |
| 259 | >332_333HK | PAMMAAAQAALQHKWGMMGML | HHHHHHHHHHHHHLLLLLLLL |
| 260 | >332_333RK | PAMMAAAQAALQRKWGMMGML | HHHHHHHHHHHHHLLLLLLLL |
| 261 | >332_333DK | PAMMAAAQAALQDKWGMMGML | HHHHHHHHHHHHHLLLLLLLL |
| 262 | >332_333TK | PAMMAAAQAALQTKWGMMGML | HHHHHHHHHHHHHLLLLLLLL |
| 263 | >332_333LM | PAMMAAAQAALQLMWGMMGML | HHHHHHHHHHHHHLLLLLLLL |
| 264 | >332_333LF | PAMMAAAQAALQLFWGMMGML | HHHHHHHHHHHHHLLLLLLLL |
| 265 | >332_333LH | PAMMAAAQAALQLHWGMMGML | HHHHHHHHHHHHHTLLLLLLL |
| 266 | >332_333HL | PAMMAAAQAALQHLWGMMGML | HHHHHHHHHHHHHLLLLLLLL |
| 267 | >332_333MW | PAMMAAAQAALQMWWGMMGML | HHHHHHHHHHHHHLLLLLLLL |
| 268 | >332_333MV | PAMMAAAQAALQMVWGMMGML | HHHHHHHHHHHHHLLLLLLLL |
| 269 | >332_333MY | PAMMAAAQAALQMYWGMMGML | HHHHHHHHHHHHHLLLLLLLL |
| 270 | >332_333MH | PAMMAAAQAALQMHWGMMGML | HHHHHHHHHHHHHTLLLLLLL |
| 271 | >332_333MD | PAMMAAAQAALQMDWGMMGML | HHHHHHHHHHHHHLLLLLLLL |
| 272 | >332_333KM | PAMMAAAQAALQKMWGMMGML | HHHHHHHHHHHHHLLLLLLLL |
| 273 | >332_333QM | PAMMAAAQAALQQMWGMMGML | HHHHHHHHHHHHHLLLLLLLL |
| 274 | >332_333EM | PAMMAAAQAALQEMWGMMGML | HHHHHHHHHHHHHLLLLLLLL |
| 275 | >332_333VM | PAMMAAAQAALQVMWGMMGML | HHHHHHHHHHHHHLLLLLLLL |
| 276 | >332_333RM | PAMMAAAQAALQRMWGMMGML | HHHHHHHHHHHHHLLLLLLLL |
| 277 | >332_333NM | PAMMAAAQAALQNMWGMMGML | HHHHHHHHHHHHHLLLLLLLL |
| 278 | >332_333EQ | PAMMAAAQAALQEQWGMMGML | HHHHHHHHHHHHHLLLLLLLL |
| 279 | >332_333VQ | PAMMAAAQAALQVQWGMMGML | HHHHHHHHHHHHHLLLLLLLL |
| 280 | >332_333IQ | PAMMAAAQAALQIQWGMMGML | HHHHHHHHHHHHHLLLLLLLL |
| 281 | >332_333HQ | PAMMAAAQAALQHQWGMMGML | HHHHHHHHHHHHHLLLLLLLL |
| 282 | >332_333RQ | PAMMAAAQAALQRQWGMMGML | HHHHHHHHHHHHHLLLLLLLL |
| 283 | >332_333NQ | PAMMAAAQAALQNQWGMMGML | HHHHHHHHHHHHHLLLLLLLL |
| 284 | >332_333DQ | PAMMAAAQAALQDQWGMMGML | HHHHHHHHHHHHHLLLLLLLL |
| 285 | >332_333QE | PAMMAAAQAALQQEWGMMGML | HHHHHHHHHHHHHLLLLLLLL |
| 286 | >332_333QY | PAMMAAAQAALQQYWGMMGML | HHHHHHHHHHHHHLLLLLLLL |
| 287 | >332_333QH | PAMMAAAQAALQQHWGMMGML | HHHHHHHHHHHHHLLLLLLLL |
| 288 | >332_333QT | PAMMAAAQAALQQTWGMMGML | HHHHHHHHHHHHHLLLLLLLL |
| 289 | >332_333NR | PAMMAAAQAALQNRWGMMGML | HHHHHHHHHHHHHLLLLLLLL |
| 290 | >332_333DR | PAMMAAAQAALQDRWGMMGML | HHHHHHHHHHHHHLLLLLLLL |
| 291 | >332_333TR | PAMMAAAQAALQTRWGMMGML | HHHHHHHHHHHHHLLLLLLLL |
| 292 | >332_333RT | PAMMAAAQAALQRTWGMMGML | HHHHHHHHHHHHHLLLLLLLL |
| 293 | >332_333HV | PAMMAAAQAALQHVWGMMGML | HHHHHHHHHHHHHLLLLLLLL |
| 294 | >332_333VY | PAMMAAAQAALQVYWGMMGML | HHHHHHHHHHHHHLLLLLLLL |
| 295 | >332_333WN | PAMMAAAQAALQWNWGMMGML | HHHHHHHHHHHHHLLLLLLLL |
| 296 | >332_333KW | PAMMAAAQAALQKWWGMMGML | HHHHHHHHHHHHHTLLLLLLL |
| 297 | >332_333NW | PAMMAAAQAALQNWWGMMGML | HHHHHHHHHHHHHLLLLLLLL |
| 298 | >332_333YY | PAMMAAAQAALQYYWGMMGML | HHHHHHHHHHHHHTLLLLLLL |
| 299 | >332_333HY | PAMMAAAQAALQHYWGMMGML | HHHHHHHHHHHHHTTLLLLLL |
| 300 | >332_333RY | PAMMAAAQAALQRYWGMMGML | HHHHHHHHHHHHHLLLLLLLL |
| 301 | >332_333YH | PAMMAAAQAALQYHWGMMGML | HHHHHHHHHHHHHTLLLLLLL |
| 302 | >332_333GD | PAMMAAAQAALQGDWGMMGML | HHHHHHHHHHHHLLLLGLLLL |
| 303 | >332_333RG | PAMMAAAQAALQRGWGMMGML | HHHHHHHHHHHHLLLLGLLLL |
| 304 | >332_333TQ | PAMMAAAQAALQTQWGMMGML | HHHHHHHHHHHHLLLLLLLLG |
| 305 | >332_333NY | PAMMAAAQAALQNYWGMMGML | HHHHHHHHHHHHTLLLGLLLL |
| 306 | >332_333DY | PAMMAAAQAALQDYWGMMGML | HHHHHHHHHHHHTLLLGLLLL |
| 307 | >332_333SN | PAMMAAAQAALQSNWGMMGML | HHHHHHHHHHHLLLLGGLLLL |
| 308 | >332_333GP | PAMMAAAQAALQGPWGMMGML | HHHHHHHHHHLLLLGLGLLLG |
| 309 | >332_333PL | PAMMAAAQAALQPLWGMMGML | HHHHHHHHHHLLLLGLLLLGG |
| 310 | >332_333Wildtype | PAMMAAAQAALQSSWGMMGML | HHHHHHHHHHHHLLLLLLLLL |
| 311 | >332_333SL | PAMMAAAQAALQSLWGMMGML | HHHHHHHHHHHHLLLLLLLLL |
| 312 | >332_333SK | PAMMAAAQAALQSKWGMMGML | HHHHHHHHHHHHLLLLLLLLL |
| 313 | >332_333SQ | PAMMAAAQAALQSQWGMMGML | HHHHHHHHHHHHLLLLLLLLL |
| 314 | >332_333SE | PAMMAAAQAALQSEWGMMGML | HHHHHHHHHHHHLLLLLLLLL |
| 315 | >332_333SS | PAMMAAAQAALQSSWGMMGML | HHHHHHHHHHHHLLLLLLLLL |
| 316 | >332_333SV | PAMMAAAQAALQSVWGMMGML | HHHHHHHHHHHHLLLLLLLLL |
| 317 | >332_333SI | PAMMAAAQAALQSIWGMMGML | HHHHHHHHHHHHLLLLLLLLL |
| 318 | >332_333SD | PAMMAAAQAALQSDWGMMGML | HHHHHHHHHHHHLLLLLLLLL |
| 319 | >332_333GS | PAMMAAAQAALQGSWGMMGML | HHHHHHHHHHHHLLLLLLLLL |
| 320 | >332_333AS | PAMMAAAQAALQASWGMMGML | HHHHHHHHHHHHLLLLLLLLL |
| 321 | >332_333LS | PAMMAAAQAALQLSWGMMGML | HHHHHHHHHHHHLLLLLLLLL |
| 322 | >332_333MS | PAMMAAAQAALQMSWGMMGML | HHHHHHHHHHHHLLLLLLLLL |
| 323 | >332_333FS | PAMMAAAQAALQFSWGMMGML | HHHHHHHHHHHHLLLLLLLLL |
| 324 | >332_333QS | PAMMAAAQAALQQSWGMMGML | HHHHHHHHHHHHLLLLLLLLL |
| 325 | >332_333SS | PAMMAAAQAALQSSWGMMGML | HHHHHHHHHHHHLLLLLLLLL |
| 326 | >332_333VS | PAMMAAAQAALQVSWGMMGML | HHHHHHHHHHHHLLLLLLLLL |
| 327 | >332_333IS | PAMMAAAQAALQISWGMMGML | HHHHHHHHHHHHLLLLLLLLL |
| 328 | >332_333CS | PAMMAAAQAALQCSWGMMGML | HHHHHHHHHHHHLLLLLLLLL |
| 329 | >332_333HS | PAMMAAAQAALQHSWGMMGML | HHHHHHHHHHHHLLLLLLLLL |
| 330 | >332_333NS | PAMMAAAQAALQNSWGMMGML | HHHHHHHHHHHHLLLLLLLLL |
| 331 | >332_333DS | PAMMAAAQAALQDSWGMMGML | HHHHHHHHHHHHLLLLLLLLL |
| 332 | >332_333DD | PAMMAAAQAALQDDWGMMGML | HHHHHHHHHHHHLLLLLLLLL |
| 333 | >332_333TD | PAMMAAAQAALQTDWGMMGML | HHHHHHHHHHHHLLLLLLLLL |
| 334 | >332_333DE | PAMMAAAQAALQDEWGMMGML | HHHHHHHHHHHHLLLLLLLLL |
| 335 | >332_333TE | PAMMAAAQAALQTEWGMMGML | HHHHHHHHHHHHLLLLLLLLL |
| 336 | >332_333EY | PAMMAAAQAALQEYWGMMGML | HHHHHHHHHHHHTTLLLLLLL |
| 337 | >332_333ED | PAMMAAAQAALQEDWGMMGML | HHHHHHHHHHHHLLLLLLLLL |
| 338 | >332_333FK | PAMMAAAQAALQFKWGMMGML | HHHHHHHHHHHHLLLLLLLLL |
| 339 | >332_333FE | PAMMAAAQAALQFEWGMMGML | HHHHHHHHHHHHLLLLLLLLL |
| 340 | >332_333FI | PAMMAAAQAALQFIWGMMGML | HHHHHHHHHHHHLLLLLLLLL |
| 341 | >332_333FT | PAMMAAAQAALQFTWGMMGML | HHHHHHHHHHHHLLLLLLLLL |
| 342 | >332_333NF | PAMMAAAQAALQNFWGMMGML | HHHHHHHHHHHHTLLLLLLLL |
| 343 | >332_333DF | PAMMAAAQAALQDFWGMMGML | HHHHHHHHHHHHTLLLLLLLL |
| 344 | >332_333GA | PAMMAAAQAALQGAWGMMGML | HHHHHHHHHHHHLLLLLLLLL |
| 345 | >332_333GK | PAMMAAAQAALQGKWGMMGML | HHHHHHHHHHHHTLLLLLLLL |
| 346 | >332_333GQ | PAMMAAAQAALQGQWGMMGML | HHHHHHHHHHHHTLLLLLLLL |
| 347 | >332_333GE | PAMMAAAQAALQGEWGMMGML | HHHHHHHHHHHHLLLLLLLLL |
| 348 | >332_333GH | PAMMAAAQAALQGHWGMMGML | HHHHHHHHHHHHLLLLLLLLL |
| 349 | >332_333GR | PAMMAAAQAALQGRWGMMGML | HHHHHHHHHHHHTLLLLLLLL |
| 350 | >332_333GN | PAMMAAAQAALQGNWGMMGML | HHHHHHHHHHHHLLLLLLLLL |
| 351 | >332_333GT | PAMMAAAQAALQGTWGMMGML | HHHHHHHHHHHHTLLLLLLLL |
| 352 | >332_333AG | PAMMAAAQAALQAGWGMMGML | HHHHHHHHHHHHLLLLLLLLL |
| 353 | >332_333HD | PAMMAAAQAALQHDWGMMGML | HHHHHHHHHHHHLLLLLLLLL |
| 354 | >332_333HI | PAMMAAAQAALQHIWGMMGML | HHHHHHHHHHHHLLLLLLLLL |
| 355 | >332_333DI | PAMMAAAQAALQDIWGMMGML | HHHHHHHHHHHHLLLLLLLLL |
| 356 | >332_333IN | PAMMAAAQAALQINWGMMGML | HHHHHHHHHHHHLLLLLLLLL |
| 357 | >332_333IT | PAMMAAAQAALQITWGMMGML | HHHHHHHHHHHHLLLLLLLLL |
| 358 | >332_333KD | PAMMAAAQAALQKDWGMMGML | HHHHHHHHHHHHLLLLLLLLL |
| 359 | >332_333QK | PAMMAAAQAALQQKWGMMGML | HHHHHHHHHHHHLLLLLLLLL |
| 360 | >332_333VK | PAMMAAAQAALQVKWGMMGML | HHHHHHHHHHHHLLLLLLLLL |
| 361 | >332_333NK | PAMMAAAQAALQNKWGMMGML | HHHHHHHHHHHHLLLLLLLLL |
| 362 | >332_333LE | PAMMAAAQAALQLEWGMMGML | HHHHHHHHHHHHLLLLLLLLL |
| 363 | >332_333LT | PAMMAAAQAALQLTWGMMGML | HHHHHHHHHHHHLLLLLLLLL |
| 364 | >332_333DL | PAMMAAAQAALQDLWGMMGML | HHHHHHHHHHHHLLLLLLLLL |
| 365 | >332_333MM | PAMMAAAQAALQMMWGMMGML | HHHHHHHHHHHHLLLLLLLLL |
| 366 | >332_333MF | PAMMAAAQAALQMFWGMMGML | HHHHHHHHHHHHLLLLLLLLL |
| 367 | >332_333MK | PAMMAAAQAALQMKWGMMGML | HHHHHHHHHHHHLLLLLLLLL |
| 368 | >332_333MQ | PAMMAAAQAALQMQWGMMGML | HHHHHHHHHHHHLLLLLLLLL |
| 369 | >332_333ME | PAMMAAAQAALQMEWGMMGML | HHHHHHHHHHHHLLLLLLLLL |
| 370 | >332_333MP | PAMMAAAQAALQMPWGMMGML | HHHHHHHHHHHHLLLLLLLLL |
| 371 | >332_333MR | PAMMAAAQAALQMRWGMMGML | HHHHHHHHHHHHLLLLLLLLL |
| 372 | >332_333MN | PAMMAAAQAALQMNWGMMGML | HHHHHHHHHHHHLLLLLLLLL |
| 373 | >332_333MT | PAMMAAAQAALQMTWGMMGML | HHHHHHHHHHHHLLLLLLLLL |
| 374 | >332_333HM | PAMMAAAQAALQHMWGMMGML | HHHHHHHHHHHHLLLLLLLLL |
| 375 | >332_333DM | PAMMAAAQAALQDMWGMMGML | HHHHHHHHHHHHLLLLLLLLL |
| 376 | >332_333TM | PAMMAAAQAALQTMWGMMGML | HHHHHHHHHHHHLLLLLLLLL |
| 377 | >332_333NN | PAMMAAAQAALQNNWGMMGML | HHHHHHHHHHHHLLLLLLLLL |
| 378 | >332_333DN | PAMMAAAQAALQDNWGMMGML | HHHHHHHHHHHHLLLLLLLLL |
| 379 | >332_333TN | PAMMAAAQAALQTNWGMMGML | HHHHHHHHHHHHLLLLLLLLL |
| 380 | >332_333NT | PAMMAAAQAALQNTWGMMGML | HHHHHHHHHHHHLLLLLLLLL |
| 381 | >332_333QQ | PAMMAAAQAALQQQWGMMGML | HHHHHHHHHHHHLLLLLLLLL |
| 382 | >332_333QV | PAMMAAAQAALQQVWGMMGML | HHHHHHHHHHHHLLLLLLLLL |
| 383 | >332_333QD | PAMMAAAQAALQQDWGMMGML | HHHHHHHHHHHHLLLLLLLLL |
| 384 | >332_333NV | PAMMAAAQAALQNVWGMMGML | HHHHHHHHHHHHLLLLLLLLL |
| 385 | >332_333DV | PAMMAAAQAALQDVWGMMGML | HHHHHHHHHHHHLLLLLLLLL |
| 386 | >332_333TV | PAMMAAAQAALQTVWGMMGML | HHHHHHHHHHHHLLLLLLLLL |
| 387 | >332_333VN | PAMMAAAQAALQVNWGMMGML | HHHHHHHHHHHHLLLLLLLLL |
| 388 | >332_333VT | PAMMAAAQAALQVTWGMMGML | HHHHHHHHHHHHLLLLLLLLL |
| 389 | >332_333GG | PAMMAAAQAALQGGWGMMGML | HHHHHHHHHHHLLLLLLLLLG |
| 390 | >332_333LG | PAMMAAAQAALQLGWGMMGML | HHHHHHHHHHHLLLLLGLLLL |
| 391 | >332_333QG | PAMMAAAQAALQQGWGMMGML | HHHHHHHHHHHLLLLLGLLLL |
| 392 | >332_333NG | PAMMAAAQAALQNGWGMMGML | HHHHHHHHHHHLLLLLGLLLL |
| 393 | >332_333PM | PAMMAAAQAALQPMWGMMGML | HHHHHHHHHLLLLLGLLLLGG |
| 394 | >332_333SG | PAMMAAAQAALQSGWGMMGML | HHHHHHHHHHHLLLLLLLLLL |
| 395 | >332_333GW | PAMMAAAQAALQGWWGMMGML | HHHHHHHHHHHLLLLLLLLLL |
| 396 | >332_333KG | PAMMAAAQAALQKGWGMMGML | HHHHHHHHHHHLLLLLLLLLL |
| 397 | >332_333EG | PAMMAAAQAALQEGWGMMGML | HHHHHHHHHHHLLLLLLLLLL |
| 398 | >332_333HG | PAMMAAAQAALQHGWGMMGML | HHHHHHHHHHHLLLLLLLLLL |
| 399 | >332_333LP | PAMMAAAQAALQLPWGMMGML | HHHHHHHHHHHTLTTLLLLLL |
| 400 | >332_333QP | PAMMAAAQAALQQPWGMMGML | HHHHHHHHHHHLLLLLLLLLL |
| 401 | >332_333NP | PAMMAAAQAALQNPWGMMGML | HHHHHHHHHHLLLLTLLLLLG |
| 402 | >332_333PH | PAMMAAAQAALQPHWGMMGML | HHHHHHHHHHLLLLLLLLLLL |
